# Missense mutation in the activation segment of the kinase CK2 models Okur Chung neurodevelopmental disorder and alters the hippocampal glutamatergic synapse

**DOI:** 10.1101/2024.04.04.588084

**Authors:** Jose M. Cruz-Gamero, Demetra Ballardin, Barbara Lecis, Chun-Lei Zhang, Laetitia Cobret, Alexander Gast, Severine Morisset-Lopez, Rebecca Piskorowski, Dominique Langui, Joachim Jose, Guillaume Chevreux, Heike Rebholz

## Abstract

Exome sequencing has enabled the identification of causative genes of monogenic forms of autism, amongst them, in 2016, CSNK2A1, the gene encoding the catalytic subunit of the kinase CK2, linking this kinase to Okur-Chung Neurodevelopmental Syndrome (OCNDS), a newly described neurodevelopmental condition with many symptoms resembling those of autism spectrum disorder.

Thus far, no preclinical model of this condition exists. Here we describe a knockin mouse model that harbors the K198R mutation in the activation segment of the kinase. This region is a mutational hotspot, representing one-third of patients. These mice exhibit behavioral phenotypes that mirror patient symptoms. Homozygous knock-in (KI) mice die mid-gestation while heterozygous KI mice are born at half of the expected mendelian ratio and are smaller in weight and size than wildtype littermates. Heterozygous KI mice showed alterations in cognition and memory-assessing paradigms, enhanced stereotypies, altered circadian activity patterns, and nesting behavior. Phosphoproteome analysis from brain tissue revealed alterations in the phosphorylation status of major pre- and postsynaptic proteins of heterozygous KI mice. In congruence, we detect reduced synaptic maturation in hippocampal neurons and attenuated long-term potentiation in the hippocampus of KI mice. Taken together, K198R KI mice exhibit significant face validity, presenting ASD-relevant phenotypes, synaptic deficits and alterations in synaptic plasticity, all of which strongly validate this line as a mouse model of OCNDS.

## Introduction

In the US, approximately 15% of children are affected by neurodevelopmental disorders (NDDs) (1), a group of conditions characterized by delayed or impaired central nervous system function and maturation, and include autism spectrum disorder (ASD), intellectual disability and a general delay in reaching developmental milestones (2), resulting from a combination of genetic and environmental risk factors (3).

In 2016, WES analysis led to the identification of mutations in the gene encoding the catalytic subunit of the kinase CK2 in the Okur-Chung neurodevelopmental syndrome (OCNDS, OMIM #617062), an ASD-type condition characterized by developmental delay, intellectual disability, behavioral problems (hyperactivity, repetitive movements and social interaction deficits), hypotonia and language/verbalization deficits (*4–11*). CK2 is a ubiquitously-expressed heterotetrameric kinase consisting of 2 catalytic CK2α subunits or α’ (12), and 2 regulatory CK2β subunits (13). CK2 is upregulated in cancers and CK2 inhibitors have shown a beneficial effect in preclinical tumor models (14, 15). Interestingly, CK2 is highly expressed in the brain (16) and CK2α^(-/-)^ embryos die mid-gestation, due to severe defects in brain and heart development (17). We have shown that CK2α is a modulator of receptor endocytosis and neurotransmitter signaling of Gα_s_-coupled receptors such as D1, A2a and 5-HT4 receptors (18, 19). Drd1a-Cre;CK2α KO mice exhibit behavioral phenotypes characterized by hyperactivity, enhanced stereotypy, hyper-exploration, reduced motor learning, nesting behavior and altered circadian rhythm (20).

OCNDS-linked CSNK2A1 mono-allelic variants can be found along the whole kinase domain and are in their majority missense mutations. Basic information on the effect of OCNDS-causing mutations on protein folding, function, kinase activity or substrate specificity is still rudimentary. Most mutations are localized in the activation loop or the ATP binding domain, strongly supporting the hypothesis that loss of function underlies symptoms. This is also suggested by deletion and splicing variants yielding no or severely truncated protein. We have shown that 15 different OCNDS-linked missense mutations, either in the form of purified GST-fusion proteins or when overexpressed and immunoprecipitated from mammalian cells, exhibit reduced kinase activity towards peptide substrates, albeit to different degrees (21). The main hotspot is K198R comprising ca. 30% of all known point mutations. We had previously found the activity of ectopically expressed and of purified recombinant CK2α K198R mutant protein towards a substrate peptide to be reduced by ca. 90% and 80%, respectively (21). X-ray crystallography has revealed a significant structural shift supporting the notion of alteration of substrate specificity (22, 23).

To understand the effect of individual mutations and identify pathological mechanisms affected in OCNDS, we characterized a novel knock-in mouse line, CK2^K198R^. Our goal in this work is to establish if this line has face validity and identify cellular functions and pathways that may underlie the condition.

## Material & Methods

A detailed description of the materials and methods is provided as supplemental file.

### Animals

CK2^K198R^ mice (C57BL/6J-Csnk2a1em2Lutzy/Mmjax) carry a K198R point mutation in the *Csnk2a1* gene (MMRRC stock #67399, The Jackson Laboratory). Animals were maintained on the C57BL/6J background. Experiments were performed in adult male mice (4-7 months), maintained in a 12 h light/dark cycle, with access to food/water *ad libitum*. Animal procedures were approved by the French Ministry of Higher Education and are in accordance with the local IACUC (CEEA-34, Apafis No. 2022011423502271).

### Statistics

All data are presented as meanL±LSEM (*pL<L0.05; **pL<L0.01; ***pL<L0.001; ****pL<L0.0001).

## Results

### Reduced viability of K198R^(+/wt)^ embryos

In this newly developed murine model, homozygosity of the K198R variant leads to embryonic death, similar to findings in CK2α KO mice (17). Thus, our study was performed using Wt and heterozygous mice (referred as K198R^(+/wt)^ or Het in the text).

K198R^(+/wt)^ mice are born at a lower-than-expected mendelian ratios. Wt:Het is reduced from the expected 1:1 to 1:0.42 in Wt x K198R^(+/wt)^ matings, and from 1:2 to 1:0.72 in K198R^(+/wt)^ x K198R^(+/wt)^ matings (fig. 1A,B). Male to female K198R^(+/wt)^ birth ratio was 1, as in the human condition. At E10.5, K198R^(+/+)^ presented with edematous heart defects (marked by arrow in fig. 1C), the blood was not circulating, and the reabsorption of the yolk sac had started. At E11.5, homozygotes were non-viable, and some showed a failure to close the cranium (fig. 1C, see arrow-head). There were viable K198R^(+/wt)^ embryos that resembled Wt, and others with heart defects (marked by arrow in fig. 1C) resembling homozygotes at E10.5. At E15.5, from a total of seven yolk sacs, 4 (2 K198R^(+/+)^ and 2 K198R^(+/wt)^), were completely reabsorbed, while two Wt and one K198R^(+/wt)^ developed correctly (fig. 1C). Post weaning, K198R^(+/wt)^ males were more prone to die in the first 18 months of age (mortality 12,4% in K198R^(+/wt)^ compared to 1.4% in Wt) while K198R^(+/wt)^ females were affected from 3-18 months (mortality 12,3% in K198R^(+/wt)^ compared to 0.8% in Wt), supporting a trend of less viability in K198R^(+/wt)^ mice (suppl. table S1A).

**Figure 1.**
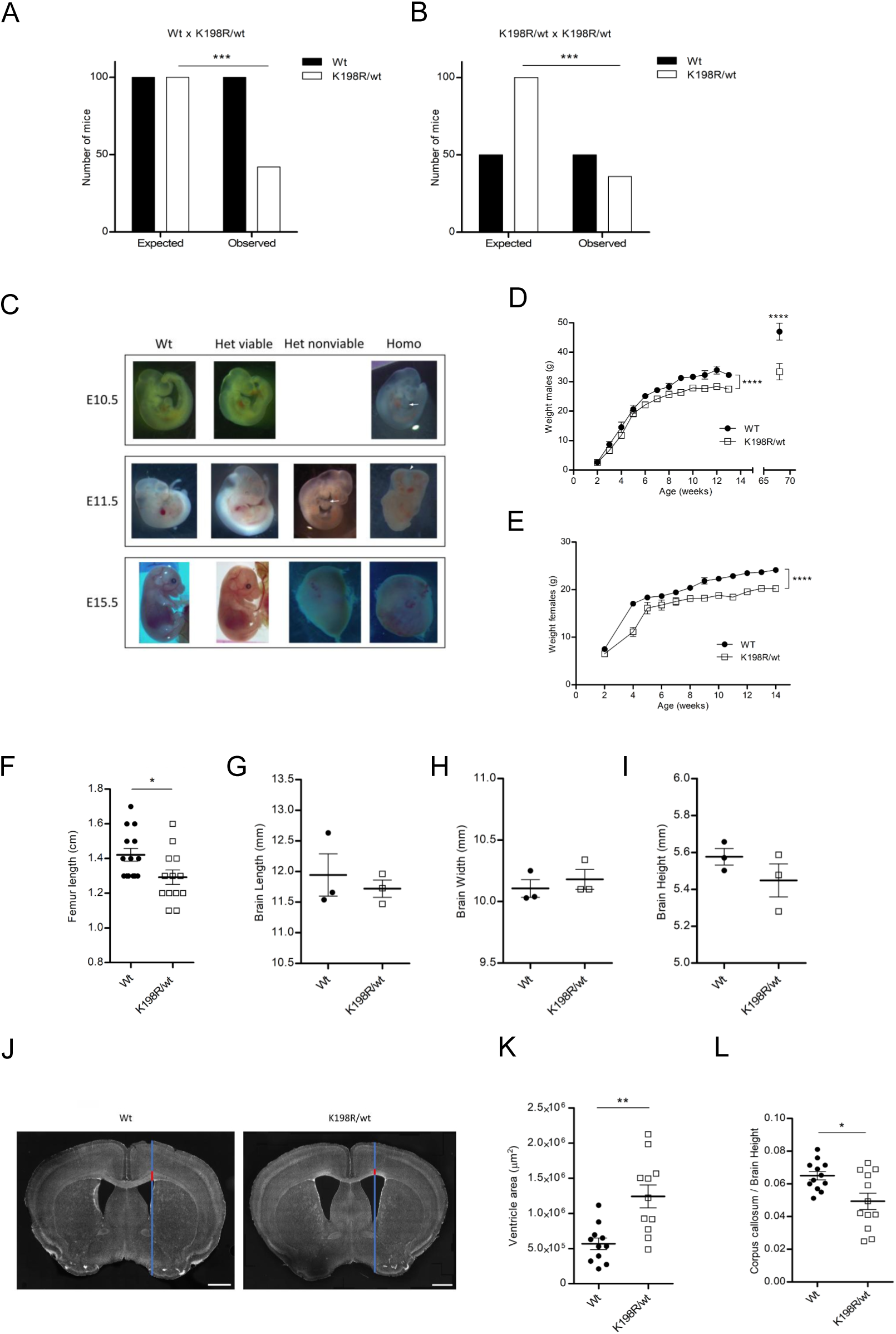
Gross phenotyping comparison of K198R^(+/wt)^ mice. In Wt × K198R^(+/wt)^ and K198R^(+/wt)^ × K198R^(+/wt)^ matings, the observed number of heterozygous born was strongly reduced compared to the expected ratios (*p* = 0.0002 and *p* = 0.0003, respectively, Fisher’s exact test) **(A,B)**. Photos of embryos at different gestational stages. Arrows point to enlarged heart. Arrowhead indicates unclosed cranial portion **(C)**. Body weight curves of Wt and K198R^(+/wt)^ males (*n* = 7 Wt and *n* = 9 Het) and females (*n* = 10 Wt and *n* = 7 Het) from birth until adulthood. Het weigh less than Wt (males: *F*(1,106) = 45.75, *p* < 0.0001 for genotype; females: *F*(1,153) = 180.4, *p* < 0.0001 for genotype; 2-way ANOVA); and the difference increases in males at 68 weeks (*p* < 0.0001, Bonferroni post-test, *n* = 7 Wt and *n* = 5 Het) **(D,E).** Femur length is reduced in K198R/wt mice (1.421 ± 0.03658 cm in Wt versus 1.292 ± 0.04154 cm in Het; *p* = 0.0275; unpaired *t* test, *n* = 14 Wt and *n* = 13 Het) **(F)**. Measurements of length, width and height of coronal brain slices show no difference between genotypes (*p* = 0.5816, 0.5324, and 0.2715 respectively; unpaired *t* test, *n* = 3 per genotype) **(G,H,I).** Hoechst-stained sections of Wt and K198R^(+/wt)^ mice (70 µm). Blue/red lines show positioning of measurements. Scale bars: 1 mm **(J)**. Lateral ventricle area measurements, Het: 1.24 ± 0.16 mm^2^, Wt: 0.57 ± 0.08 mm^2^; *p* = 0.0014; unpaired *t* test, *n* = 6 mice/genotype) **(K)**. Corpus callosum height normalized by total brain height: 0.049 ± 0.005 (Het) and 0.065 ± 0.002 (Wt), *p* = 0.0105; unpaired *t* test, *n* = 6 mice/genotype **(L)**.

### K198R^(+/wt)^ mice weigh less and show gross brain alterations

The majority of OCNDS patients exhibit growth problems such as microcephaly and reduced weight and stature (24). We observed that male and female K198R mice weighed 16.07% and 17.6% less, respectively, at 14 weeks of age, a difference that increases with age for males reaching nearly 33.9% at 16 months (fig. 1D,E). K198R^(+/wt)^ mice are smaller, as assessed per femur length (fig. 1F, suppl. fig. S1B). The weight difference cannot be explained by a difference in food intake which is not altered (suppl. fig. S1C). Since CK2 is necessary for neural tube development (8), we analyzed whether gross brain architecture was impacted in K198R^(+/wt)^ mice. No obvious alterations in brain size (length, width, height) or gross structure were detected (fig. 1G-I), however, the lateral ventricle was enlarged (fig. 1J,K), while the thickness of the corpus callosum was reduced (fig. 1J,L), a finding reminiscent of observations in ASD patients and other preclinical models (25, 26).

### K198R variant leads to diminished CK2**α** expression and activity

Western blotting using striatal tissue showed that expression of the CK2α, α’ and β subunits were unaltered in K198R^(+/wt)^ mice compared to Wt (fig. 2A-D). qPCR confirmed that CK2α mRNA expression was similarly equal between genotypes (fig. 2E). Subcellular localization of CK2α was determined by immunohistochemical analysis using coronal brain slices. No changes were detected, CK2α being predominantly cytosolic and excluded from nucleoli (suppl. fig. S2). Since we had previously shown that recombinant CK2αK198R protein exhibits reduced kinase activity (13), we next analyzed the activity of brain extracts from Het and Wt mice towards a substrate peptide. We detected 25% reduction of CK2 activity in K198R^(+/wt)^ mice (fig. 2F and suppl. fig. S3). Consequently, we determined if phosphorylation levels of bona fide CK2 substrates, such as AKT and DARPP-32 were affected. For pDARPP-32, we observed a significant decrease in phosphorylation of the CK2 site, S97 (fig. 2G,H) while the phosphorylation in Thr34 (PKA site) was unchanged (fig. 2G,I). pS129 AKT, shown to be a direct CK2 substrate *in vitro* and in cancer cells (27) was not altered (fig. 2G,J), confirming prior data from patient fibroblasts (21). We further measured elevated phosphorylation of S473 Akt, a PDK1 substrate and an indicator of the activation state of Akt (fig. 2G,K). These findings are interesting since a dysregulated mTor/AKT pathway is a recurring finding in ASD (*28*). We further tested S6 Ribosomal Protein S240/244 phosphorylation, a downstream readout of the mTor pathway, and found it increased in heterozygotes (fig. 2G,L). Altogether, these experiments indicate that K198R variant affects kinase activity but not subunit stability or subcellular localization.

**Figure 2.**
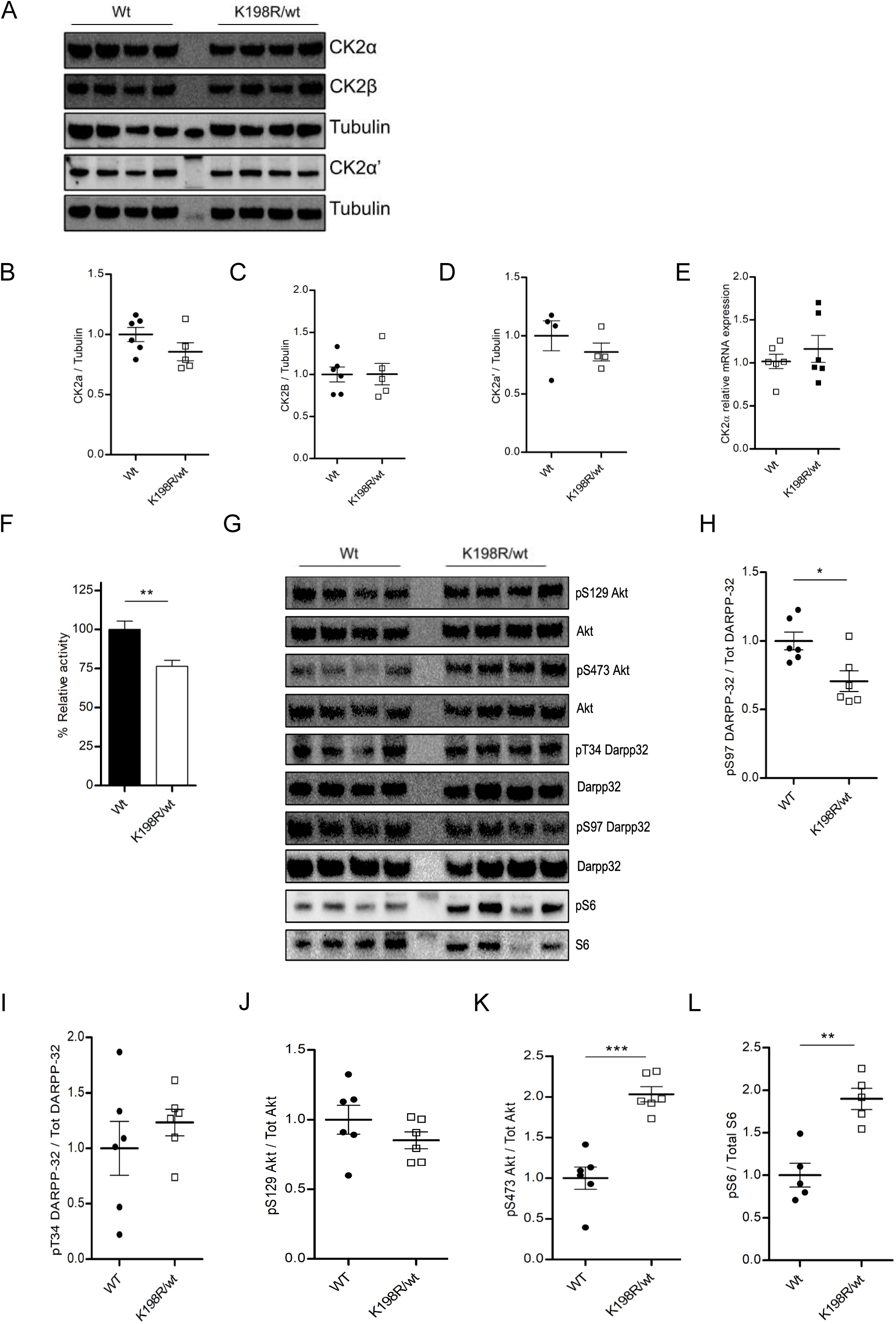
Expression and activity of CK2 in K198R(+/wt) mice. Western blot analysis of striatal lysates to assess CK2 subunits levels **(A)** and quantification thereof showed no differences in protein levels (*p* = 0.1615, 0.9795 and 0.3879, respectively) **(B-D)**. qPCR using probes show no difference in relative mRNA expression of CK2a (*p* = 0.4327) **(E)**. CK2 activity was measured by capillary electrophoresis-based separation showing reduced activity in K198R(+/wt) (*p* = 0.0031; unpaired *t* test, *n* = 3/genotype) **(F)**. Western blot analysis to assess phosphorylation levels of CK2 substrates **(G).** pS97 DARPP-32 is reduced (*p* = 0.0143), but not pT34 DARPP-32 (*p* = 0.4077) **(H,I)** or pS129 AKT (*p* = 0.2458) **(J),** while pS473 AKT (*p* = 0.0001) **(K)** and pS240/244 S6 Ribosomal Protein are increased (*p* = 0.0013) **(L)**. Unless otherwise stated, *n* was 4-6/genotype, unpaired *t* test was used.

We wanted to determine if the interaction of mutant CK2α with the regulatory β subunit is altered *in vivo*, which could impact substrate specificity as the beta subunit is known to mediate recruitment of certain substrates (29). To this end, Bioluminescence Resonance Energy Transfer (BRET) analysis was performed in HEK-293 cells, transiently transfected with expression vectors for CK2bβ-NanoLuciferase and CK2Wt-eYFP or CK2K198R-eYFP (suppl fig. S4). BRET saturation assays were developed to monitor, quantify, and compare the formation of CK2 heteromers. We did not observe any changes in the BRETmax (which represents the relative amounts of dimers) or the BRET50 value (which represents the relative dimer affinity) for K198R mutant protein (suppl fig. S4B-D).

### Mass spectrometry

To detect overall changes in the phosphoproteome, we performed mass spectrometric analysis of striatal tissue which revealed 13017 unique phosphopeptides (p-peptides), out of which 394 were significantly altered by a factor of 1.5 or more between genotypes. 163 were present at higher levels in Wt striatum, while 231 peptides were enriched in K198R^(+/wt)^ mice (fig. 3A). As depicted in the volcano plot and heatmap, phosphopeptides from synaptic proteins or synaptic vesicle transport proteins (e.g. SYNAPSIN1, SynGAP1, SGIP1) and membrane- and cytoskeleton-associated proteins, e.g. pTau or pMAP1A were identified. In K198R^(+/wt)^ males, we found upregulation of phosphopeptides of the presynaptic protein Piccolo, which has been linked to synaptic vesicle recycling (30), as well as Grk6 and several MARKs (fig. 3B,C). Gene Ontology analysis pointed to the synapse as the most affected cellular compartment and the glutamatergic synapse was also identified in this search (fig. 3D). We used Metascape to create protein association networks, whereby the 5 top networks involved synaptic interaction networks such as the Neurexin, neuregulin network, a generic phosphorylation network including SynGAP1 and GSK3 (fig. 3E). In conclusion, multiple bioinformatic analyses converged on the synapse.

**Figure 3.**
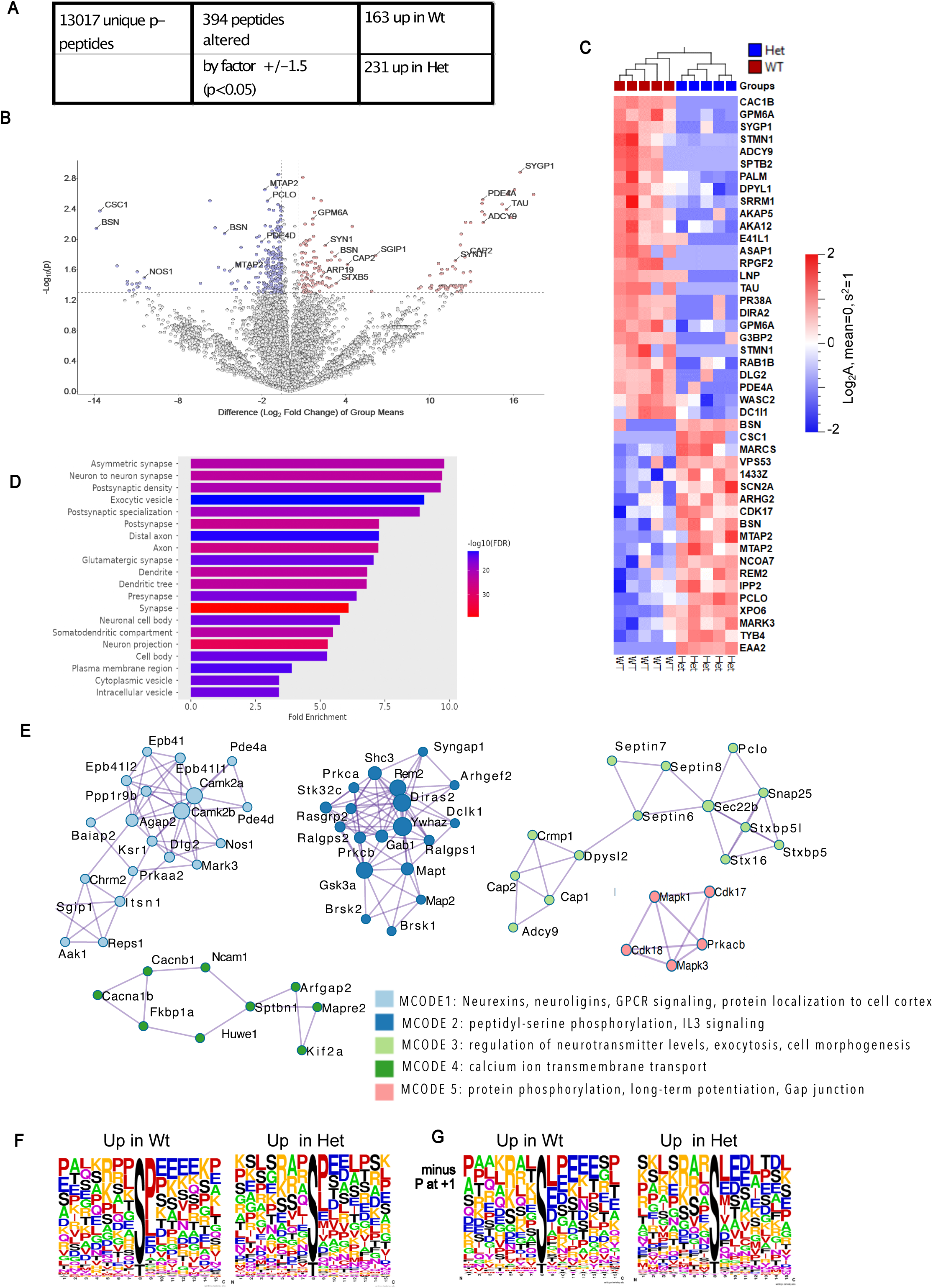
Mass spectrometric analysis of phosphoproteome. Table depicting the unique enriched phosphopeptides (*n* = 5/genotype), cutoff for fold change of ± 1.5 and *p*-value < 0.05 **(A).** Volcano plot depicting identified peptides, with cutoff and *p*-values as in A. **(B)**. Heatmap showing gene names corresponding to altered p-proteins in 5 Wt and 5 Het mouse striata using a more stringent cutoff for fold change of ± 2 and *p*-value < 0.01, to simplify the pictogram **(C)**. Gene ontology analysis for cellular compartment using a cutoff for fold change of ± 1.5 and *p*-value < 0.05 **(D)**. Protein-protein interaction enrichment analysis: 5 top network clusters that were most significantly altered when comparing Wt and Het are depicted **(E)**. Depiction of peptide sequence weblogos **(F),** we eliminated p-peptides with proline at position +1 **(G)**.

To assess the potential contribution of other kinases that may be regulated by CK2, we generated weblogos containing 7 amino acids prior and after the phosphosite. The minimal consensus sequence recognized by CK2 is (S/T-x-x-E/D/pX) (31). We found that the majority of peptides reduced in Het (“Up in Wt’) have a proline at position +1, this was much less the case the peptides that are reduced in Wt, even though proline and leucine are still the most prominent amino acids at position +1. An acidic patch or 4 glutamates (but not aspartates) appears in all altered p-peptides, but was present to a higher extent in the Wt. The difference in the presence of these acidic residues may be attributable to loss of CK2 activity (fig. 3F). Interestingly, in the Het we detected more p-peptides with aspartate at position +3 (fig. 3F). When peptides containing proline at +1 are excluded, the acidic residues at +3 to +4 are enhanced. In this panel, it is interesting that at +3, representing the crucial acidic residue for recognition by CK2, the percentage is slightly higher in the peptides that are more phosphorylated in Het, indicating that mutant CK2 K198R may directly be involved in phosphorylation of these peptides (fig. 3G).

### K198R variant causes defects in hippocampal synapses

Since hippocampal volume is affected in ASD patients (32) and many preclinical ASD models present defects in hippocampal function (33, 34) and since GO analysis of our MS results pointed towards alteration at the glutamatergic synapse, we examined the hippocampus of K198R mice by electron microscopy. Neither number of presynaptic vesicles per synapse nor thickness of the postsynaptic density was changed (fig. 4A, B), however, the disc diameter was significantly enhanced in Het animals (fig. 4C). Furthermore, synapse density was enhanced in K198R^(+/wt)^, while overall cell number in the same region is not changed (fig. 4D,E). Representative images of glutamatergic synapses are shown in figure 4F.

**Figure 4.**
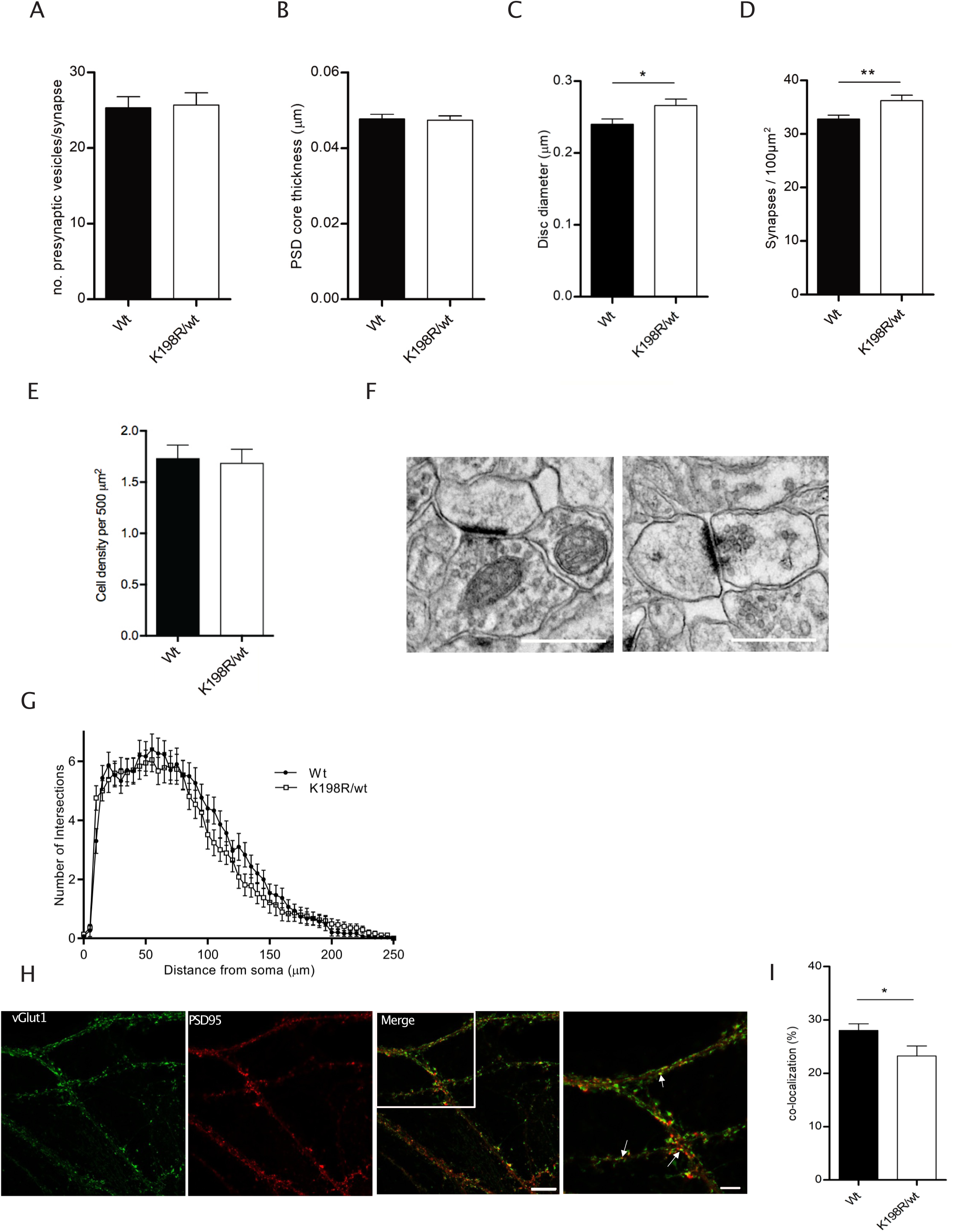
Synapse analysis in hippocampus. 99 images of 35.62 µm^2^ size (between 23-49 images per mouse, *n* = 3 /genotype) were analyzed by electron microscopy. Number of presynaptic vesicles (*p* = 0.9941) **(A)**. Postsynaptic density core thickness (*p* = 0.7329) **(B)**; PSD length (0.24 ± 0.01 µm in Wt, 0.27 ± 0.01 µm in Het; *p* = 0.0252) **(C)** and Synapse density (32.9 ± 0.77 in Wt versus 36.4 ± 1.05 synapses/µm^2^ in K198R^(+/wt)^ mice; *p* = 0.0058) **(D)** were plotted. Cell density is based on DAPI counts in a confocal image of 0.54 mm^2^ of the CA1 region (*p* = 0.8072, *n* = 6/genotype) **(E)**. Representative EM micrographs of CA1 regions, at a resolution of 494 pixels/µm **(F)**. Statistical analysis was performed using unpaired *t* test. In primary hippocampal culture, no differences between genotypes were detected by Sholl analysis (*p* = 0.5442 for Genotype; 2-way ANOVA, *n* = 37 cells from 7 of mice for Wt, *n* = 30 cells from 6 mice for K198R^(+/wt)^) **(G)**. Immunostaining of hippocampal neurons using anti-PSD95 and anti-vGLUT1 antibodies **(H)** and quantification of the juxtaposition of signals in % using anti-VGLUT1 and PSD-95 antibodies (28.03 ± 1.25% in Wt versus 23.24 ± 1.89% in K198R^(+/wt)^; *p* = 0.0369; unpaired *t* test; *n* = 18 images/6 Wt mice and *n* = 15 images/ 5 K198R^(+/wt)^ mice) **(I)**. Scale bars = 10 µm.

In cultured hippocampal primary neurons, we performed the 2D Sholl analysis to determine extent of dendritic branching. When we compared Wt and K198R^(+/wt)^ intersections, we found no differences in the dendritic arbor (fig.4G). We then analyzed the composition of the synapses formed after 14 days in culture, by visualizing the presynaptic glutamate transporter VGLUT1 and the postsynaptic density scaffold, PSD-95. We counted the number of VGLUT1+ and PSD-95+ puncta and determined the percentage of juxtaposed signals derived from both antibodies, as a proxy for the synapse formation. We found that in K198R^(+/wt)^ hippocampal cultures, synapse formation is reduced compared to Wt (fig. 4H,I).

These alterations suggest that the K198R variant affects the formation of glutamatergic synapses. We postulated that while synaptic transmission itself is likely functional, synaptic plasticity may be altered. To test this, we performed field recordings in acute hippocampal slices from Wt and K198R^(+/wt)^ mice. We recorded excitatory postsynaptic potentials (fEPSPs) in area CA1 by stimulating the Schaffer collateral inputs from area CA3 (fig. 5A,B). Using a theta-burst stimulation (TBS) induction protocol, we observed normal induction and expression of long-term potentiation (LTP) in the hippocampal slices prepared from Wt mice, but an attenuation of LTP in slices from K198R^(+/wt)^ mice (fig. 5A-C, fig. 5D shows corresponding fiber volleys). LTP in K198R^(+/wt)^ showed a significant reduction post-induction as well as a lower increase in the early expression phase of LTP (15 min) and even more so during the later phase of LTP (fig. 5C,E). Thus, these results suggested that the CK2αK198R variant produces an effect on attenuating the synaptic plasticity in the hippocampus, which may underlie certain behavioral deficits.

**Figure 5.**
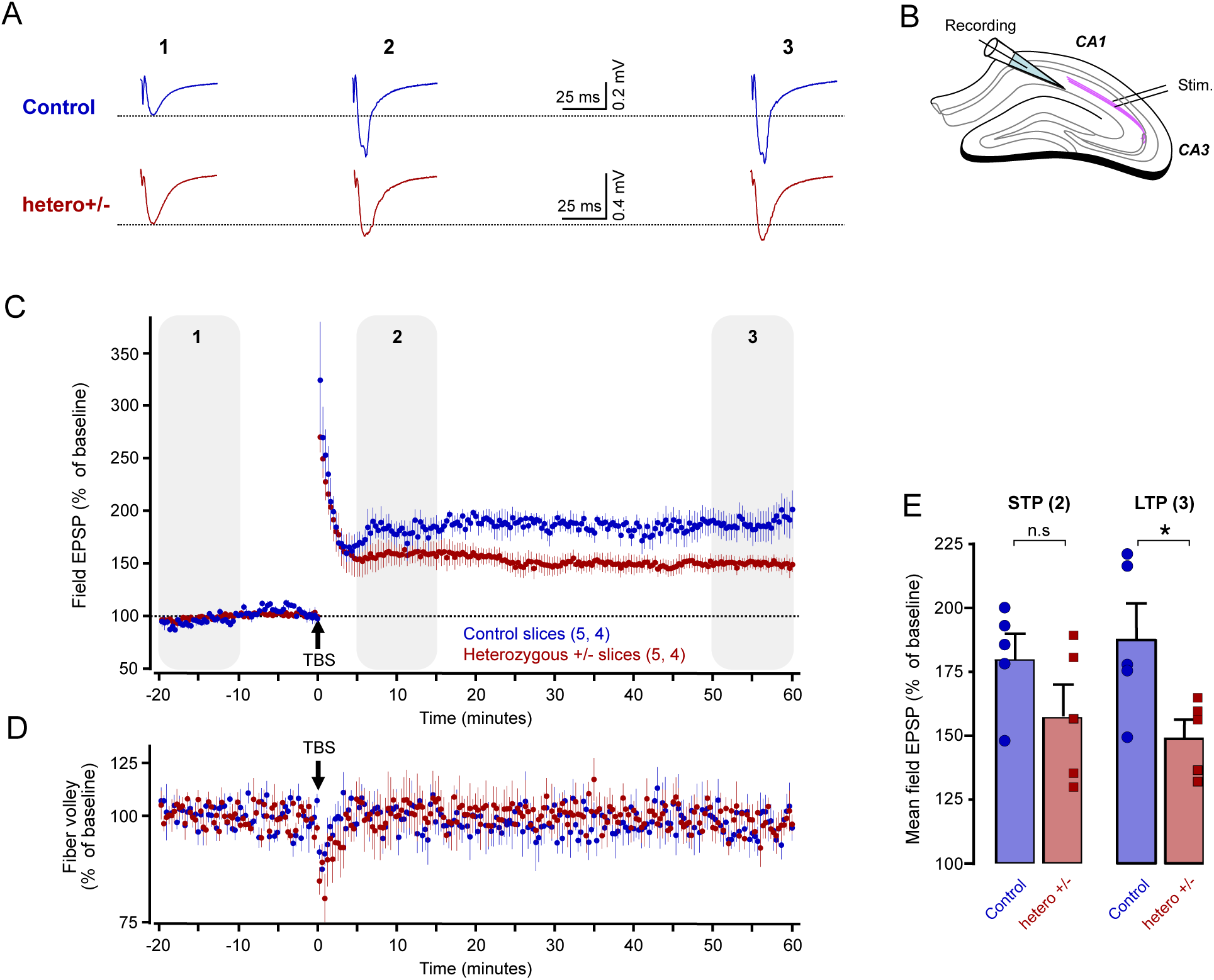
Synaptic potentiation of hippocampal CA1 field EPSPs *in vitro*. Sample traces of field excitatory post synaptic potentials (fEPSPs) for Wt (blue) and K198R^(+/wt)^ (red) at three representative points during long-term potentiation (LTP), 1: at the baseline period, 2: the early-phase of the LTP, and 3: the late-phase of the LTP. **(A)**. Synaptic transmission was evoked by stimulation of CA3 Schaffer collateral fibers and recorded with a pipette in the CA1 stratum radiatum (SR) **(B)**. Summary graph shows the time course of normalized fEPSPs, before and after theta-burst stimulation (TBS; *n* = 5 slices from 4 mice per genotype) **(C)**. Normalized fiber volley for both genotypes **(D)**. Graph showing the short-term and long-term LTP amounts. At 5-15 minutes following the TBS, at the early phase of LTP expression there is no significant difference between the genotypes, whereas a significant attenuation is observed in K198R^(+/wt)^ slices at 50-60 minutes post TBS, the long-term phase (short term phase: *p* = 0.9769; long term phase: *p* = 0.0332). Statistical analysis was performed using unpaired *t* test.

### K198R^(+/wt)^ mice show defects in memory-related behaviors

We next wanted to assess if the K198R mutation would impact mouse behavior. We tested exploratory behavior and working memory in the Y-maze spontaneous alternation test and found that K198R^(+/wt)^ mice performed fewer correct alternations compared to Wt (fig. 6A), while in the Y-maze recognition test, there was no difference between genotypes (suppl. fig. S5A). We examined spatial learning performance in the Barnes maze. Both groups were able to correctly learn the location of the hidden box and made successively fewer error nose pokes during learning (suppl. fig. S5B-E) and reversal learning phases (suppl. fig. S5F-I). Although K198R^(+/wt)^ mice had shorter escape latency in the learning phases (suppl. fig. S5D,H), this seemed to be due to them having a higher velocity and not to better learning *i.e.*, same number of errors (suppl. fig. S5B,F). During the learning probe test, there was no difference between genotypes in the time spent in the zone where the escape box previously was (target zone) (fig. 6B). In the reversal learning probe test, both Wt and K198R^(+/wt)^ mice spent more time in the zone where the hidden box was during the reversal learning (new target zone) (fig. 6C). When we focused on the time spent in the zone where the escape box was during the learning phase, even though there is a trend of K198R^(+/wt)^ spending more time, no significant differences were found (fig. 6C). Thus, in the Barnes test, no deficits in spatial learning nor cognitive flexibility in heterozygotes mice was detected.

**Figure 6.**
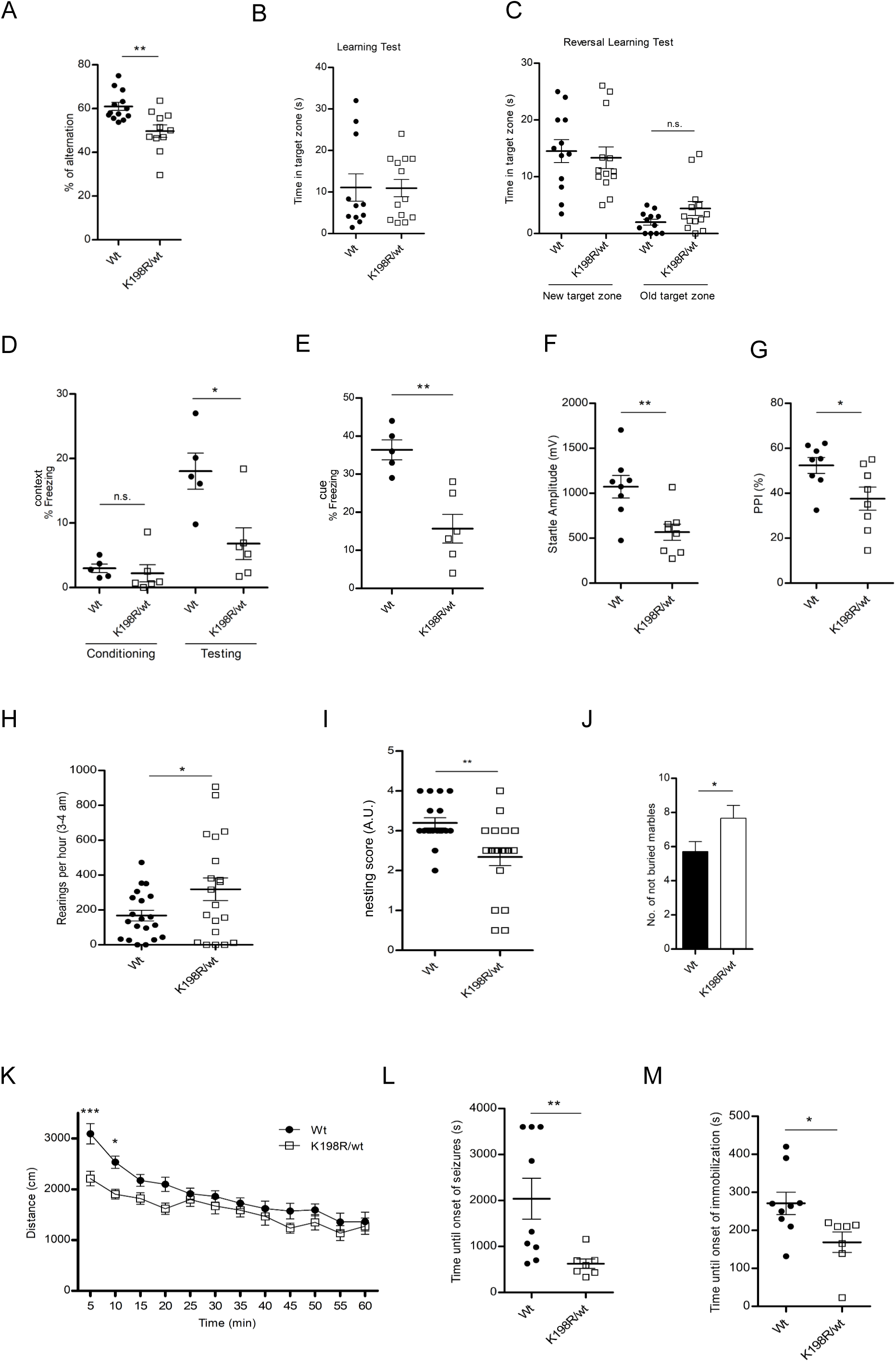
Behavioral characterization. Y-maze alternation test depicts correct alternations in % (*p* = 0.0059; *n* = 13 Wt and *n* = 11 K198R^(+/wt)^ mice) **(A)**. Time spent in the target zone during Barnes maze learning probe test (*p* = 0.9673, *n* = 11 Wt and *n* = 13 K198R^(+/wt)^) **(B)**. Time spent in the new (*p* = 0.5831) or old target zones during Barnes maze reversal test (*p* = 0.1629, *n* = 12 Wt and *n* = 13 K198R^(+/wt)^) **(C)**. Contextual Fear conditioning showed no genotypic difference during conditioning (*p* = 0.1775, *n* = 5 Wt and 6 K198R^(+/wt)^), but during testing: K198R^(+/wt)^ mice freeze less than Wt (*p* = 0.0303) **(D).** Cued freezing is significantly lower in K198R^(+/wt)^ mice than Wt (*p* = 0.0043) **(E)**. Pre-pulse inhibition (PPI) showed reduced startle amplitude in K198R^(+/wt)^ (*p* = 0.007, *n* = 8 per genotype) **(F)** and percentage of PPI compared to Wt (*p* = 0.0281) **(G)**. For all tests, statistical analysis was performed using Mann-Whitney test. Number of rearings of singly housed animals from 3-4 am after 34 hours of habituation (*p* = 0.0424; unpaired *t* test, *n* = 20/genotype) **(H)**. Nesting using cotton pads was scored after 16 hrs (overnight) (*p* = 0.001; Mann-Whitney test, *n* = 18 Wt and 19 K198R^(+/wt)^) **(I)**. The number of unburied marbles after a 30 min test is higher in K198R^(+/wt)^ mice (*p* = 0.045; unpaired *t* test, *n* = 12/genotype) **(J)**. Locomotor activity and thigmotaxis were measured in the open field environment (F(1,24) = 5.926, *p* = 0.0227 for genotype; 2-way ANOVA) (**K**). Picrotoxin (3 mg/kg; i.p.)-induced seizures were observed for one hour, time until seizure (*p* = 0.0195, Mann-Whitney test, *n* = 9 Wt and *n* = 7 K198R^(+/wt)^) **(L)** and time until immobility onset were measured (*p* = 0.0092, Mann-Whitney test, *n* = 9 Wt and *n* = 7 K198R^(+/wt)^) **(M)**. Data are presented as meanL±LSEM (*pL<L0.05; **pL<L0.01; ***pL<L0.001; ****pL<L0.0001).

When tested in the context fear conditioning paradigm, K198R^(+/wt)^ mice showed reduced fear expression 24 hours after training, suggesting an impairment in contextual fear memory compared with Wt control (fig. 6D). We also observed a lower level of cue fear expression 24 hours after training (fig. 6E). In the PPI test, a paradigm assessing sensorimotor-gating (21), K198R^(+/wt)^ mice showed reduced startle response, both expressed as reduction of startle amplitude and percentage of pre-pulse inhibition and startle amplitude (fig. 6F,G). Taken together, our results show that K198R^(+/wt)^ mice have deficits in differential behaviors related to learning and memory, congruous with the synaptic deficits observed from the electrophysiological and morphological data.

### Other ASD-relevant behavioral phenotypes

Stereotypies or restricted/repetitive behaviors are a hallmark of ASD (35). K198R^(+/wt)^ mice display enhanced stereotypies (rearings) which were assessed for 1 hr during the dark active cycle (3-4 am) cycle (fig. 6H), with a similar result obtained during the light cycle (suppl. fig. S6A). Mice also exhibited a reduced nesting score (fig. 6I). In the marble burying test (at 30 min), a test used to assess repetitive and perseverative behavior as well as anxiety, K198R^(+/wt)^ mice showed impaired marble burying behavior compared to Wt mice (fig. 6J). Mice did not exhibit overall hyperactivity, in fact in their first exposure to the open field environment, they are slightly hypoactive in the first 20 min of exposure as compared to Wt (fig. 6K).

Reduced sociability is a hallmark of ASD, and can be reproduced in mice, albeit not all mouse models of ASD present a sociability defect. To assess social behavior, we employed the 3-chamber assay. Both Wt and K198R^(+/wt)^ mice showed enhanced interest in a mouse versus an empty box, and enhanced interest towards a novel stranger mouse compared to an “already-known” mouse in the second phase. This test was performed with a cohort of mice where genotypes were mixed within cages (suppl fig. S6B) and a cohort separated by genotype at weaning, both times with no difference detected. We also monitored the behavior of pairs of males that had not been exposed to each other previously, in an open field environment. Neither number nor duration of nose-to-nose or nose-to-body contacts were altered (suppl. fig. S6C-D).

We also aimed to ascertain if mice have motor deficiencies and enhanced seizure proneness, both comorbid in OCNDS patients. In the rotarod, using the accelerated protocol over 3 consecutive days, no differences were detected between Wt and Het mice (suppl. fig. S6E). Similarly, grip strength was measured in the two-paw grip and the inverted screen test and was not affected (suppl. fig. S6F,G).

We used picrotoxin, a GABA-A antagonist to induce acute myoclonic seizures. Interestingly, K198R^(+/wt)^ mice show an earlier onset of the seizure-preceding immobilization and seizures themselves (fig. 6L,M).

Regarding mood phenotypes, we observed reduced anxiety in K198R^(+/wt)^ mice in the light dark test, and novelty suppressed feeding (suppl. fig. S7A,B). However, other, possibly more stressful paradigms, such as the elevated plus maze, tail suspension or forced swim test failed to show genotypic differences (suppl. fig. S7C-E).

## Discussion

### 1) ASD-typical abnormalities in the brain of K198R/wt mice

We found that over 50% of heterozygous K198R mice were embryonically lethal. Homozygous embryos died at E11.5, while heterozygous embryos either died between E12.5 and 15.5 or survived without a gross developmental phenotype. CK2α^(-/-)^ embryos die mid-gestation at E11.5 due to alterations in the neural tube and heart defects (17). Our findings indicate that the bi-allelic K198R mutation has a similarly detrimental impact. The high loss of K198R^(+/wt)^ embryos indicates that CK2 activity during development is tightly regulated, and that below-threshold activity leads to mortality.

K198R^(+/wt)^ mice have reduced body weight, a common finding amongst mouse models of ASD (36). We did not detect changes in brain size, or cortical thickness in adult mice. However, larger lateral ventricles in K198R^(+/wt)^ mice replicate earlier findings in ASD patients (37). Indeed, the risk of ASD diagnosis rises almost seven-fold with ventricular enlargement (38). Concomitantly with ventricle enlargement, we detect a thinning of the corpus callosum, a myelin-rich structure important for interhemispheric connectivity that has already been described to be affected in ASD patients (39).

### 2) Effect of K198R on CK2 expression, localization and activity

We did not observe a change in subcellular localization of CK2α in K198R^(+/wt)^ mice, a finding that corresponds to earlier findings in K198R patient-derived fibroblasts (21). We found *ex vivo* kinase activity of mouse brain tissue reduced by ca. 25%. Using recombinant CK2α and CK2 immunoprecipitates from cell lysates overexpressing mutant CK2α, we had previously detected a 75% and 90 % reduction in activity (21). The 25% reduction is based on overall CK2 activity, without information on which and to what extent CK2α dimers (wt-mut, wt-wt, mut-mut) participate in this activity. Based on the fact that CK2α levels in striatal lysate of K198R^(+/wt)^ mice were unaltered, we hypothesize that roughly 50% of total CK2α is comprised of mutant CK2α.

By Western blotting and MS analysis, we showed that there is a decreased phosphorylation of CK2 substrate S97 DARPP-32 (40), a prerequisite for its nuclear translocation (41). Another reported CK2 substrate, pS129 Akt, was not affected in the K198R mice, mirroring findings from patient fibroblasts (21). Since S129 Akt phosphorylation by CK2 was demonstrated in immortalized Jurkat and HEK293T cell lines (27), we hypothesize that this phosphorylation event is either cell-type specific or linked to immortalization. In contrast, the mTor pathway, known to be deregulated in ASD patients and several preclinical ASD models, such as a Shank3 insertion mutation model (42–45), is affected in K198R mice. We used S473 Akt, a downstream target of mTORC2, and pS240/244 ribosomal protein S6, a major substrate of mTORC1 (46, 47), as readouts and found both proteins to be hyperphosphorylated in K198R^(+/wt)^ mice, consistent with mTOR pathway activation participating in behavioral and synaptic phenotypes.

### 3) Phosphoproteome analysis points towards synaptic alterations

Thus, the fact that circa 3% of phosphopeptides were altered in their detection between genotypes seems appropriate. The elevated number of phosphopeptides in the brains of heterozygous mice can be explained by enhanced expression and activity of kinases that are dependent on CK2 activity. All phosphopeptides which are reduced/enhanced in K198R^(+/wt)^ animals were analyzed using weblogo and interestingly, the most prominent amino acid in the altered phosphopeptides at position n+1 is proline, followed by lysine and the two acidic residues, indicating a correlation of activity between CK2 and proline-directed kinases, as shown previously (48). Alternatively, other pleiotropic kinases which have a partial substrate consensus overlap will adopt their activity in a compensatory manner, leading to an enrichment in certain phosphopeptides. Weblogo plots show an enrichment of acidic residues at positions spanning n+2 to n+5. Aspartates were present but to a far lower extent than glutamate. The fact that the sequence surrounding the phosphorylation sites that are upregulated in K198R^(+/wt)^ mice still corresponds to the CK2 consensus (acidic residues at sites n+2 and n+3) (fig. 3F) could be interpreted either in favor of kinases with an overlapping consensus, or by altered CK2 specificity, as suggested by X-ray crystallography (22).

### 4) Synaptic deficits in OCNDS model mice

One central hallmark of ASD are synaptic deficits (49). Our study confirms such a deficit in OCNDS mice as shown by mass spectrometry. Furthermore, CK2 is expected to play a role at synapses since it was shown to co-localize with PSD-95 in primary hippocampal neurons (50). CK2 is also important for the stabilization of the pre-synapse of motor neurons (51). Finally, CK2 activity is enriched in synaptosomes (16). Electron microscopy reveals that in the CA1, the number of glutamatergic synapses per surface area is enhanced in K198R^(+/wt)^ mice, concomitantly with an enlarged diameter of the postsynaptic density. In primary hippocampal neurons, we find reduced juxtaposition of postsynaptic PSD-95 and the presynaptic glutamate transporter VGLUT1 which indicates that *in vitro,* K198R^(+/wt)^ neuronal synapse formation is attenuated, as compared to Wt neurons. This incongruence between the results from electron microscopy where more synapses per surface were counted with an enhanced thickness of the PSD disc in adult hippocampi, and the reduction of juxtaposition of VGLUT1 and PSD-95 in *in vitro* cultures was unexpected. An obvious difference between the two experiments is that the EM pictures were taken from adult mice, whereas confocal (VGLUT1-PSD-95) imaging was performed at DIV14.

CK2 inhibitors TBB and DRB were previously shown to reduce LTP at the CA3-CA1 excitatory synapse following a high-frequency stimulation; this effect was mediated through the action of CK2 on NMDA receptors (52). Interestingly, the reduction in the LTP amplitude previously reported in slices pre-incubated by a high concentration of inhibitor (100 µM) during tetanic stimulation was similar to what we observed in this work with the K198R^(+/wt)^ mice. In our experimental system, net CK2 kinase activity towards a substrate peptide is reduced to ca. 75%. We postulate that *in vivo*, or at least specifically at the CA3-CA1 glutamatergic synapse, the mutant K198R variant acts in a dominant negative manner to reduce the kinase capacity, thereby influencing synaptic plasticity. To establish this important question, further experiments would need to be performed, such as comparing slices from a CK2α^(+/-)^ mice which acts in a haploinsufficient manner.

### 5) Face validity of the novel OCNDS mouse model

Thus far, no preclinical model for the novel neurodevelopmental syndrome OCNDS has been described. We aimed to ascertain the face validity of CK2α K198R^(+/wt)^ mice. It was surprising that 50% of K198R^(+/wt)^ mice die during embryogenesis, hinting already at a robust phenotype for the surviving 50%. Cognitive deficits are apparent in the Y-maze spontaneous alternation test, without a change in Y-maze spatial recognition test, pointing towards reduction in spatial working memory. Indeed, Y-maze alternation and recognition tests reveal different entities of spatial memory: the Y-maze alternation probes mainly for an intact working memory, which requires a multi-circuitry basis from prefrontal cortex, hippocampus, septum, and basal forebrain (53, 54) while Y-maze “recognition” tests for spatial reference memory, which is strongly supported by the hippocampus (53).

The Barnes maze evaluates spatial learning and spatial memory, mainly driven by hippocampus function, during the learning phase and the learning probe trial (55, 56), while reversal learning evaluates cognitive flexibility associated with frontal cortex function (57). K198R^(+/wt)^ mice show a non-significant trend towards more time near the previous hiding place in the reversal learning phase. In this context, it is interesting that K198R^(+/wt)^ mice present reduced spatial working memory in the Y-maze alternation test, and also a trend towards reduced cognitive flexibility in the reversal phase of the Barnes paradigm, with both results pointing towards frontal cortex or dysfunctional cortical-hippocampal communication.

In the fear conditioning paradigm, heterozygous mice show reduced freezing when tested for contextual memory which is primarily hippocampal dependent, but also when cued by a tone, a type of memory that is hippocampus-independent (58). Finally, PPI associated somatosensory processing is also reduced, which has been shown to involve many brain regions including the brain stem and cortico-limbic areas like nucleus accumbens, ventral pallidum, amygdala, hippocampus, thalamus, and medial prefrontal cortex (59).

Two additional ASD core symptoms are uncontrollable repetitive movements, i.e., stereotypies, and sociability defects. We observed enhanced stereotypies (rearings in homecage) in K198R^(+/wt)^ mice. We found reduced nesting behavior and marble burying in K198R^(+/wt)^ mice. The nesting score was also reduced in the Drd1a-Cre;CK2αKO mice (19), thus confirming a role of CK2 in this phenotype. The marble burying test can be interpreted as a readout of anxiogenic effects of novelty and of stereotypy (60). Since K198R^(+/wt)^ mice show an elevated number of rearings per minute, one would also have expected an enhanced number of buried marbles - which we did not detect. However, marble burying can also be interpreted, like nesting behavior, as an index for perseverative digging behavior (61). In other models of ASDs, the marble burying assay has yielded contradicting results: for example, the number of buried marbles was also reduced in Shank1 KO mice (62), in a mouse model of Angelman’s disease (63) while in others, marble burying was enhanced such as in BTBR *T^+^tf*/J (64), Slitrk5 KO (65) or Fmr1-null mice, a model of fragile X syndrome (66). With such complex behavioral traits, we conclude that bidirectional abnormal behaviors in the marble burying paradigm can exist within different ASD models.

We found no defect in sociability or social memory in heterozygous K198R mice. For OCNDS patients, reduced sociability has not explicitly been described. Our results are consistent with numerous mouse models of ASD that do exhibit no or little sociability deficits, such as NL1 KO, Shank3 KO (Δ9) and the Shank1 KO mice (67–69).

## Conclusion

Kinases play an important role in ASDs, as attested by over 20 high-confidence kinase genes associated with ASD subtypes (gene.sfari.org). CK2 has been added to the list of kinases linked to OCNDS in 2016, and the list of clinical reports has been extended regularly. We present the K198R^(+/wt)^ KI mouse model as the first preclinical OCNDS model that exhibits construct and face validity, with a robust behavioral phenotype and plasticity deficiencies at the hippocampal glutamatergic synapse.

## Acknowledgments

We thank David Boulet, Deniz Erden, Ludivine Therreau, Gwenaelle LePen, Virginie Tolle, the animal facility team, the NeurImag facility for help and technical support. Imaging was carried out at the NeurImag facility (Inserm 1266 and Université de Paris). We thank Sésame Région Ile-de-France for funding the Zeiss880 Confocal/Airyscan system. This work was supported by the CSNK2A1 foundation (HR,JJ), by a MSCA-fellowship (894207, HR), French Ministry of Research BioSPC (JMCG, DB), Fondation pour la Recherche Médicale EQU202003010457 (RP) and Agence Nationale de la Recherche ANR-18-CE370020-01 (RP).

---

**Suppl. fig. S1.**
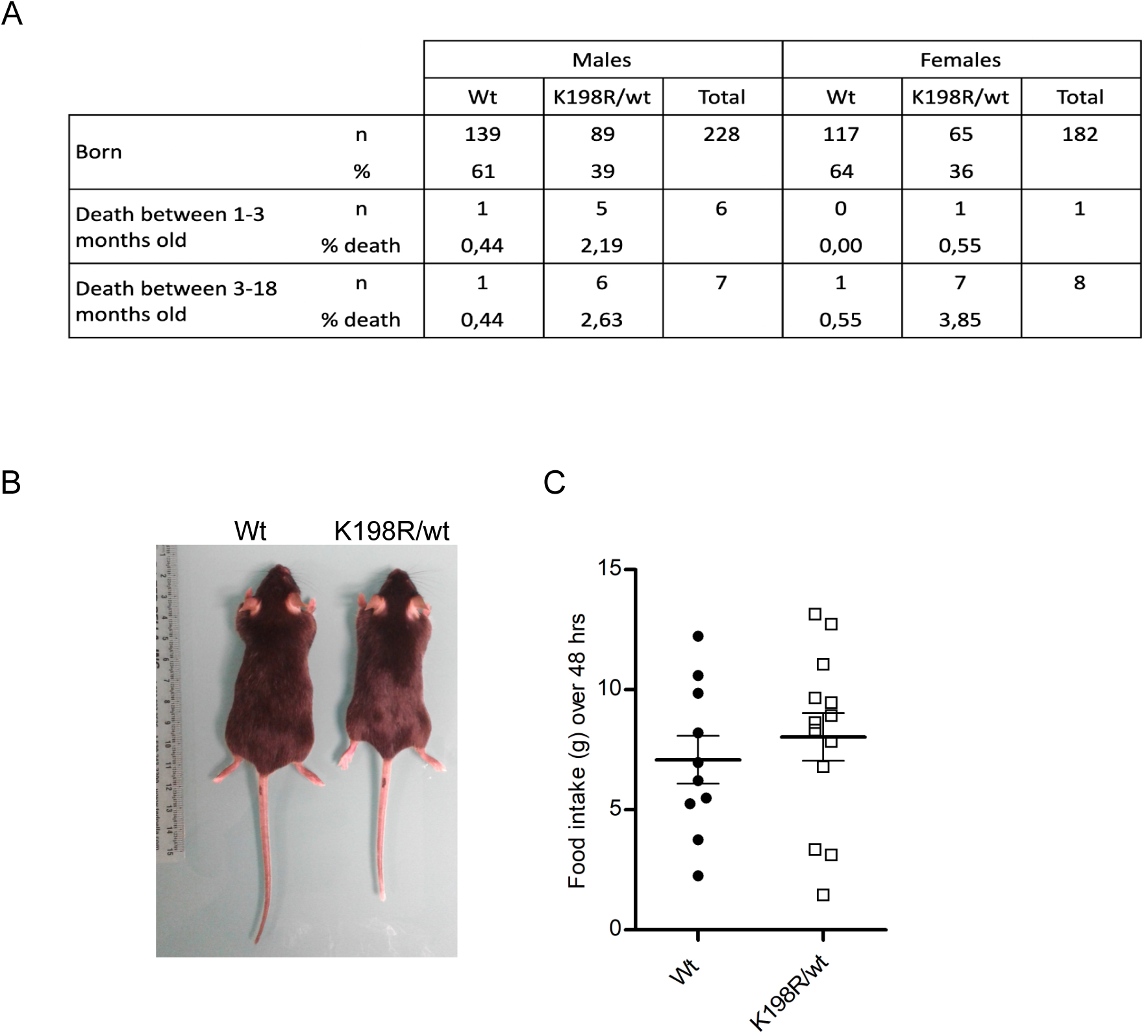

**Suppl. fig. S2.**
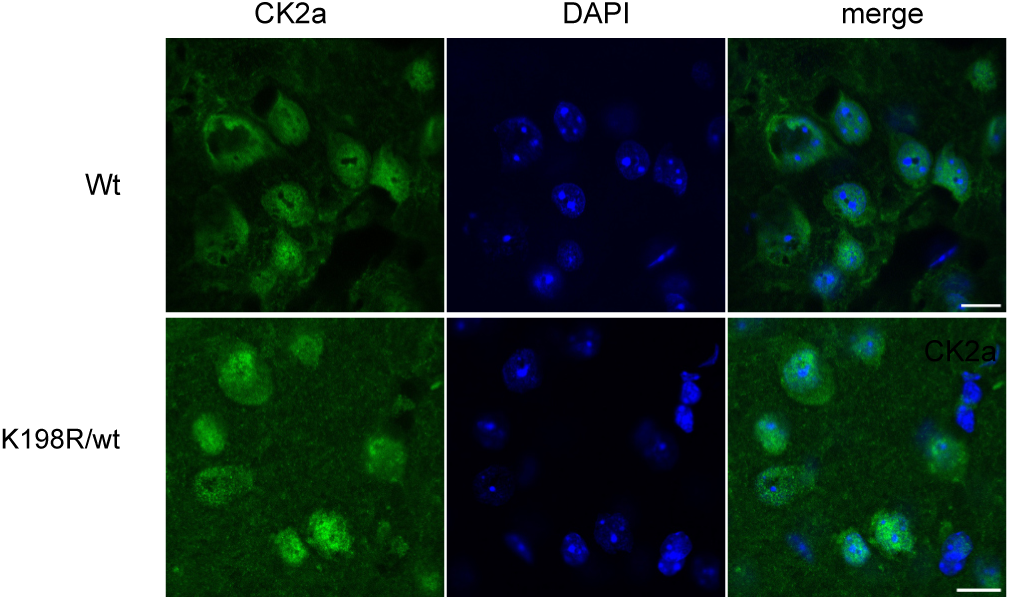

**Suppl. fig. S3.**
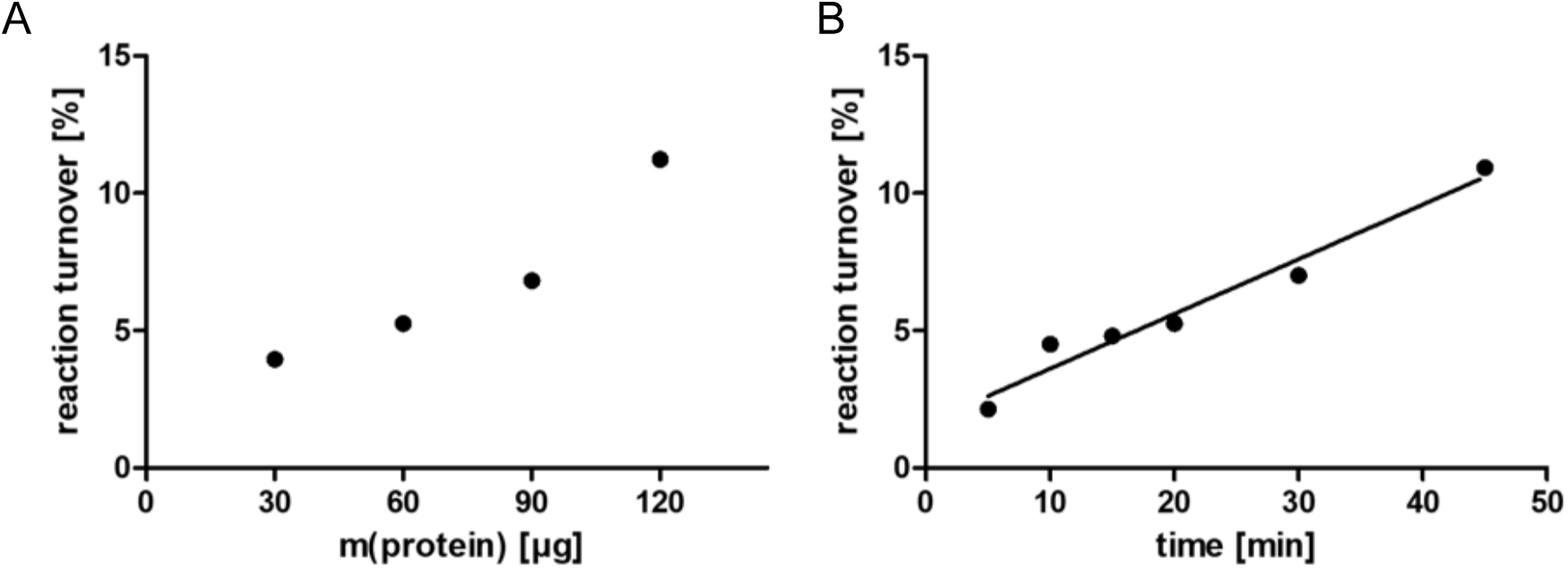

**Suppl. fig. S4.**
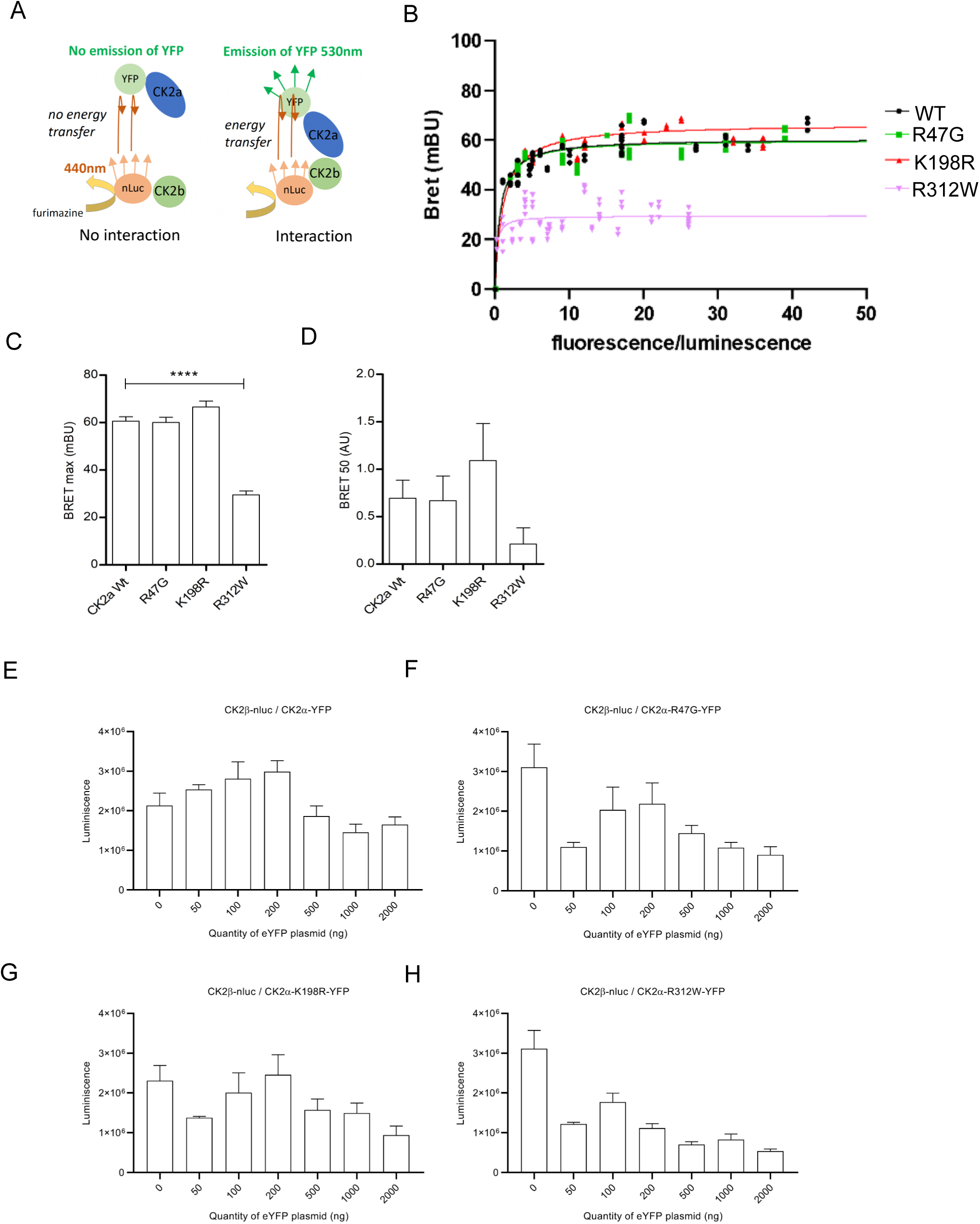

**Figure S5.**
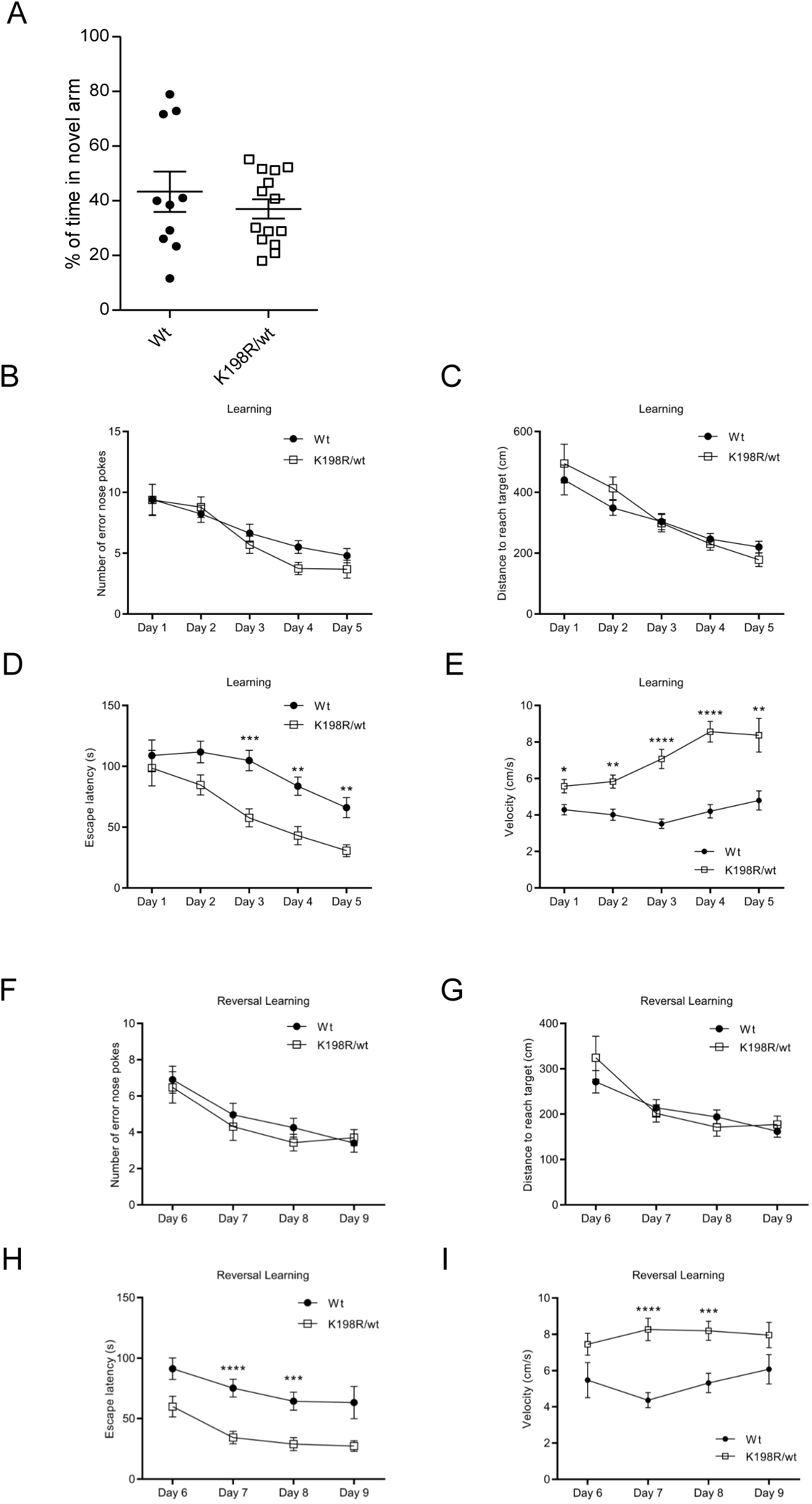
Learning and memory paradigms. Y-maze test recognition shows no differences in the time that mice spent in the novel arm (*p* = 0.7474; Mann-Whitney test, *n* = 10 Wt and *n* = 14 Het) **(A)**. Barnes maze test learning curves shows no differences between genotypes (*n* = 18 Wt and *n* = 14 K198R^(+/wt)^) for the number of error nose pokes (*F*(1,94) = 1.555, *p* = 0.2154 for Genotype) **(B)**, nor the distance travelled to reach the target (*F*(1,94) = 0.2745, *p* = 0.6016 for Genotype) **(C)**, but the escape latency was reduced in K198R^(+/wt)^ (*F*(1,94) = 17.86, *p* < 0.0001 for Genotype) **(D)**, and the velocity increased for Het compared to Wt mice (*F*(1,94) = 45.68, *p* < 0.0001 for Genotype) **(E)**. Barnes maze test reversal learning curves shows no differences between genotypes (*n* = 18 Wt and 14 K198R^(+/wt)^) for the number of error nose pokes (*F*(1,93) = 0.5002, *p* = 0.4812 for Genotype) **(F)**, nor the distance travelled to reach the target (*F*(1,93) = 0.2113, *p* = 0.6468 for Genotype) **(G)**, but the escape latency is reduced in K198R/wt mice (*F*(1,93) = 20.47, *p* < 0.0001 for Genotype) **(H)**. The velocity is increased in Het compared to Wt mice (*F*(1,93) = 14.19, *p* = 0.0003 for Genotype **(I)**. For Barnes test, statistical analysis was performed using mixed ANOVA (Bonferroni multiple comparison tests) and data are presented as mean ± SEM (*p < 0.05; **p < 0.01; ***p < 0.001; ****p < 0.0001).

**Suppl. fig. S6.**
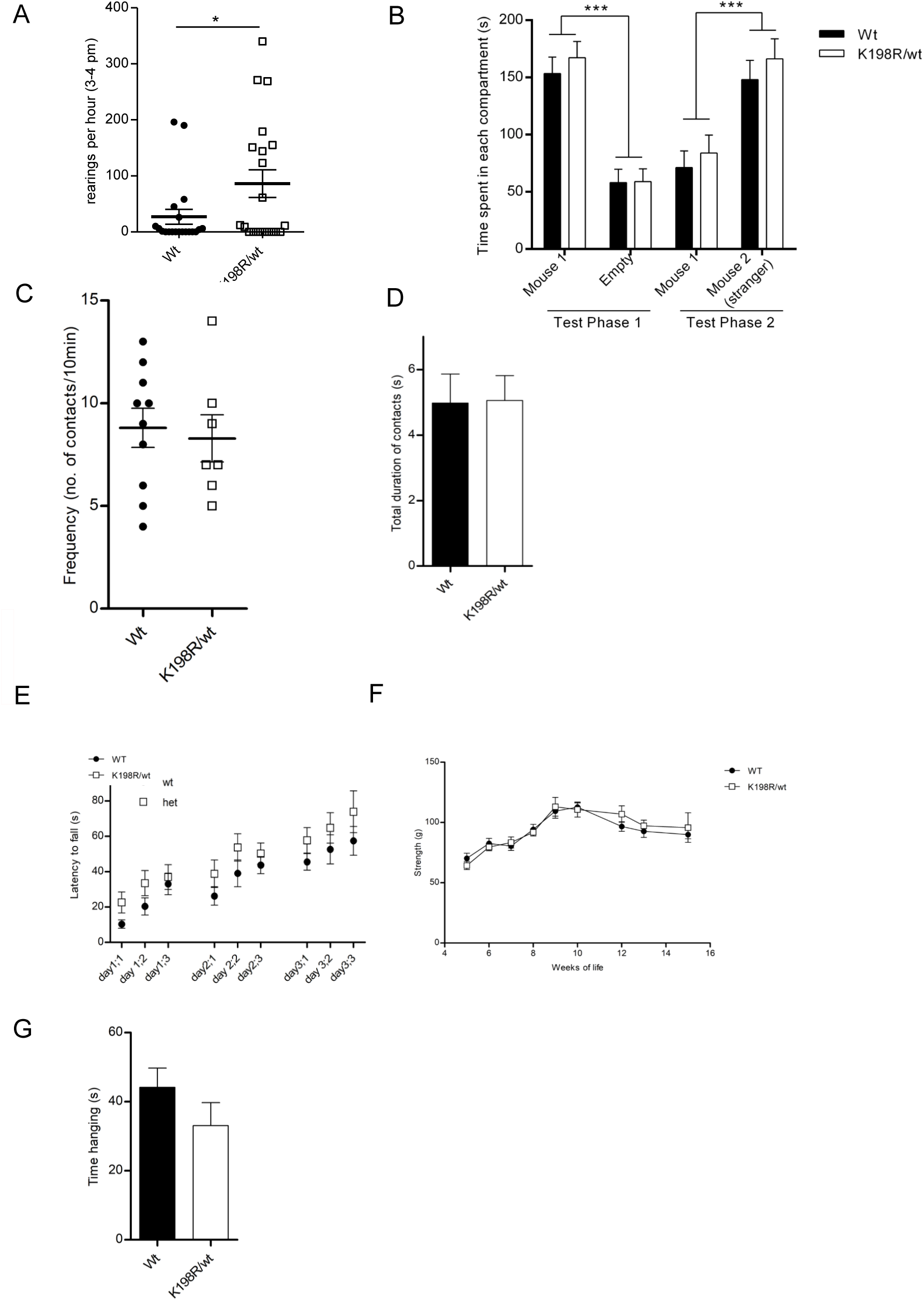

**Suppl. fig. S7.**
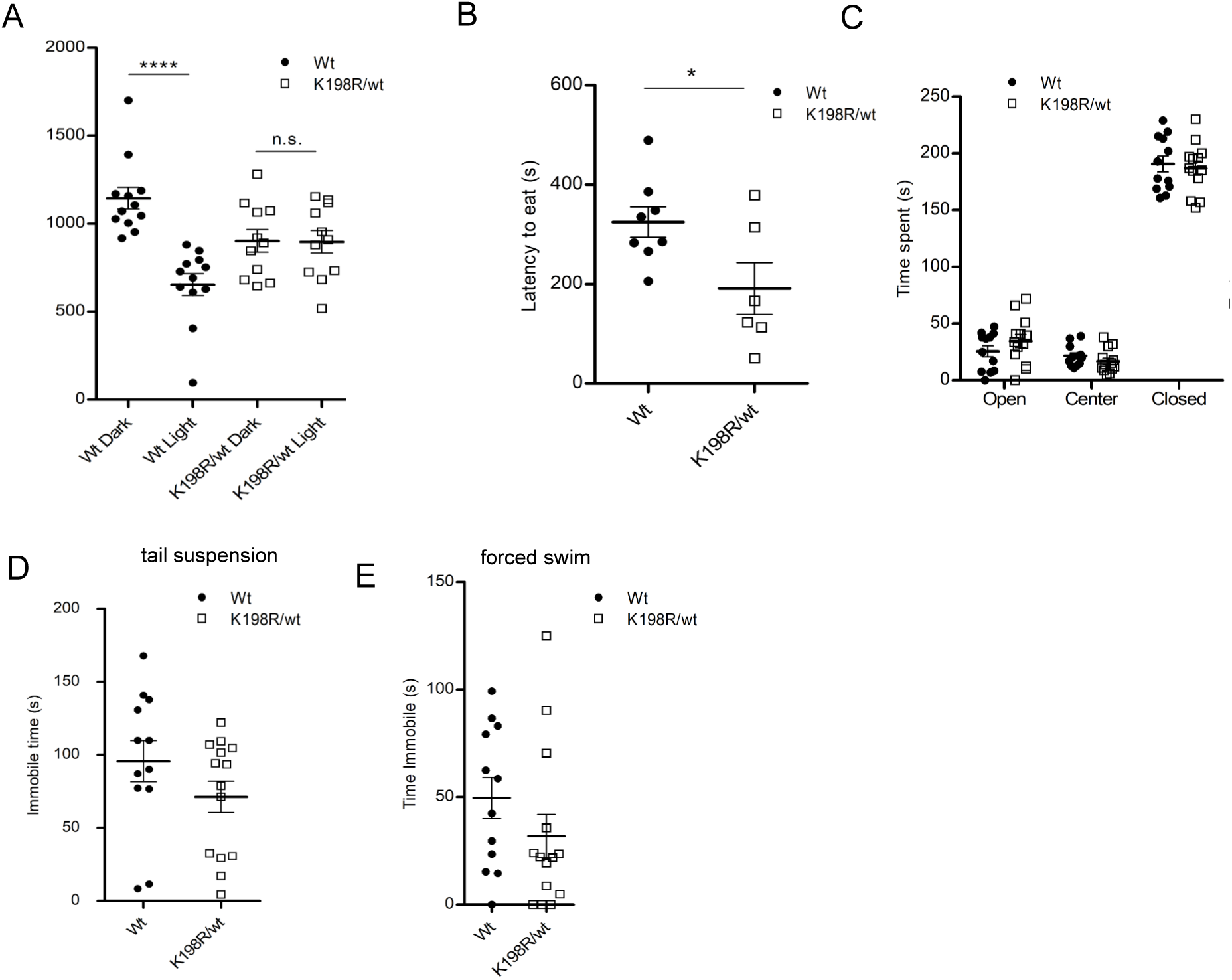

