## supplemental text for "Missense mutation in the activation segment of the kinase CK2 models Okur Chung neurodevelopmental disorder and alters the hippocampal glutamatergic synapse"

**Supplemental file**

**Methods from main text file**

**Embryo isolation**

Heterozygous K198R^(+/wt)^ mice were interbred for timed matings, and the females were checked for vaginal plugs the next morning which was considered 0.5 days after conception and thus embryonic day 0.5 (E0.5). The pregnant females were kept in their breeding cages and sacrificed for embryo collection after the appropriate number of days. Embryos were removed from the uterine horns, washed and kept in cold PBS for observation and image acquisition with a Leica M80 connected to a camera.

**Brain harvesting and dissection**

For protein or phosphorylation analyses, animals were sacrificed by decapitation and rapidly dissected after immersion of skull in liquid N_2_ to preserve phosphosites. Tissues were snap-frozen in liquid N_2_.

For perfusion, mice were anesthetized with ketamine (180 mg/kg, i.p.) / xylazine (10 mg/kg, i.p.), then intracardially perfused with 4% paraformaldehyde (PFA), and the brain was extracted and kept in 4% PFA-PBS overnight at 4 °C. The next day, brains were transferred in a 30% sucrose-PBS solution until sinking before being embedded in OCT Embedding Medium (Thermo Scientific) and stored at −80 °C. Brains were sliced in 40 μm sections with a Cryostat (Leica) and slices were stored in a cryoprotectant buffer (0.2 M PBS, 30% ethylene glycol, 30% glycerol) at −20 °C.

**Immunohistochemistry**

Perfused slices were incubated with Hoechst (Molecular probes) for nuclei staining for 30 min at RT. After 5 PBS washes, slices were mounted using VectaMount (Vector Laboratories). Images were acquired with a ZEISS Widefield Scanner inverted microscope with EC Epiplan 20x/0.4 H M27 objective without digital zoom. Widefield images consist of multiple tile regions and composite images were generated by the Zen stitching algorithm. Images were acquired with a 16-bit resolution of 1800 × 1800 pixels bidirectional laser line average of 1. Measurements of different brain regions were performed with QuPath open-source software (version 0.4.3).

**Electron microscopy**

Adult mice were deeply anesthetized by i.p. injection of ketamine (180mg/kg) / xylazine (10 mg/kg), then intracardially perfused with 2.5% glutaraldehyde and 2% paraformaldehyde in 0.1 M sodium cacodylate buffer pH 7.4 (Caco buffer). After dissection, brains were left overnight in the fixative at 4 °C. After rinsing in Caco buffer, brains were sliced in 1 mm thick coronal sections using a mouse brain acrylic matrix. Section n°6 (hippocampus level corresponding to Bregma -1.58mm) was selected, washed, post-fixed with 2% osmium tetroxide in Caco buffer for 1 h at 4 °C. After 3 washes with distilled water, sections were incubated overnight in 5% aqueous uranyl acetate, dehydrated in a graded series of ethanol solutions of 30%, 50%, 70%, 80%, 90%, and 100% (X3). Final dehydration was performed twice in 100% acetone. Samples were then progressively infiltrated with epoxy resin, Epon 812® (EMS, Souffelweyersheim, France), embedded in molds and resin polymerized at 56°C for 48 h.

Blocks were cut with an UC7 ultramicrotome (Leica Microsystems, Nanterre, France). Semi-thin sections (0.5 μm thick) were stained with 1% toluidine blue in 1% borax. Ultra-thin sections (70 nm thick) of the CA1 hippocampus region were recovered on copper grids (200 mesh, EMS, Souffelweyersheim, France) and contrasted with Reynold’s lead citrate (1). Ultrathin sections were observed with a Hitachi HT7700 electron microscope (Milexia, France) operating at 100 kV. Pictures (2048 x 2048 pixels) were taken with an AMT41B camera. Pictures were processed with ImageJ, NIH, Bethesda, USA.

**Protein extraction and Western Blot analysis**

Tissue was lysed in 50 mM Tris-HCl, pH 7.5, 150 mM NaCl, 1% NP-40, 5 mM EDTA, 1 mM EGTA, Protease inhibitor cocktail (Roche) and Phosphatase inhibitor cocktail (Roche) at 4 °C. Lysates were then centrifuged at 19 000 x g for 15 min at 4 °C and supernatants were collected. The determination of protein concentration was done with the bicinchoninic acid (BCA) protein assay kit (Thermo Fisher Scientific). Samples were denatured with 4X Sample Buffer (NuPAGE, Invitrogen). Equal amounts of protein were separated on SDS-PAGE 4-12% Bis-Tris gels (Millipore) at a constant voltage of 180 V and electro-transferred to nitrocellulose membranes (Amersham) for 50 min at 100 V. Membranes were blocked with 3% BSA in PBST for 1 hour at RT, incubated with primary antibodies overnight at 4 °C. After three washes with PBST of 5 min, membranes were incubated with secondary HRP-conjugated antibodies. When antibodies against phosphorylation sites were used, membranes were stripped in stripping buffer (62.5 mM Tris-HCl, pH 6.7, 2% SDS, 100 mM 2-mercaptoethanol) for 30 min and shaking at 50 °C, and blocked again before incubation with primary antibodies against the total protein. Bands were developed by chemiluminescence using Western Blotting Luminol reagent (Santa Cruz) and visualized by the ChemiDoc system (Bio-Rad). Quantitative analysis was performed using ImageJ software. Antibodies: rabbit anti-CK2α (A300-198A, Bethyl Laboratories, 1:2000), rabbit anti-CK2β (ab76025, Abcam, 1:1000), rabbit anti-CK2α’ (ab10474, Abcam, 1:1000), rabbit anti-pS129 AKT (5508-1, Epitomics, 1:400), rabbit anti-pS473 AKT (#4060, Cell Signaling, 1:400), mouse anti-AKT (#2920, Cell Signaling, 1:2000), rabbit anti-pS6 Ribosomal protein (Ser240/244) (#5364, Cell Signaling, 1:1000), rabbit anti-S6 Ribosomal Protein (#2217, Cell Signaling, 1:1000), rabbit anti-pT34 DARPP-32 (#12438, Cell Signaling, 1:1000), rabbit anti-pS97 DARPP-32 (#3401, Cell Signaling, 1:1000), mouse anti-DARPP-32 (sc-271111, Santa Cruz, 1:500), mouse anti-tubulin (ab44928, Abcam, 1:1000), goat anti-mouse HRP-conjugated (sc-516102, Santa Cruz, 1:10 000) and mouse anti-rabbit HRP-conjugated (sc-2357, Santa Cruz, 1:10 000).

**CK2 activity of extracts from mice brain tissue**

Tissue was homogenized in 50 mM Tris/HCl, 150 mM NaCl, 0.5% sodium deoxycholate, 1 mM NaVO4, 0.5 mM NaF, 0.8 μM aprotinin, 10 μM pepstatin A, 20 μM leupeptin, 0.1 mM PMSF, 1 mM benzamidine and 1% Triton X100 by repeated flushing through a 20G cannula. This was followed by incubation on ice for 10 min. After centrifugation (20 000 x g, 30 min, 4 °C), the supernatants were collected, and protein concentration determined. Intracellular CK2-activities were measured based on the CK2-driven phosphorylation of a fluorescently labeled peptide FITC-RRRDDDSDDD-NH_2_ (2), the reaction time was altered to 30 minutes. Quantification of conversion was conducted using a capillary electrophoresis-based separation coupled with a laser-induced fluorescence detector ^(3)^. Three activity measurements were conducted within the brain extract of each animal. Activities in wild-type mice brain extracts were set as 100%.

**Mass spectrometry**

Striatal samples were also prepared for mass spectrometry (MS) analysis in collaboration with ProteoSeine at Institut Jacques Monod, Paris, France. Tissues were lysed in RIPA buffer with Protease and Phosphatase inhibitor cocktails (Roche). A fraction was run in an SDS PAGE 4-12% Bis-Tris gel (Millipore) and a Coomassie staining was performed to check protein concentration. The samples were digested, peptides desalted, and phospho-peptides (p-peptides) enriched (Fe-NTA). Two LC-MS/MS analyses were performed, one for the proteome and another for the phosphoproteome. After trypsin digestion, the full proteome analysis was conducted on an Orbitrap Fusion mass spectrometer (ThermoFisher Scientific, Waltham, MA, USA) coupled with an Evosep one system (Evosep, Odense, Denmark) operating in long gradient mode to optimize peak capacity, resolution and sensitivity in a relatively short cycle time (48 min, SPD30 method). Phosphopeptides were enriched using the High-Select™ Fe-NTA phosphopeptide enrichment kit from ThermoFisher Scientific according to manufacturer’s procedure. Phosphopeptides were analyzed on a Q-Exactive Plus coupled to a Nano-LC Proxeon 1000, both from ThermoFisher Scientific. Mass spectrometers operated in data dependent acquisition in both experiments. MS raw files were processed using PEAKS Online X (build 1.8, Bioinformatics Solutions Inc.). Data were searched against the Mus musculus SwissProt database (04_2022, total entries 17,114). Peptide abundance was normalized on the total ion current to consider small variation of sample amount after injection. P-peptides were quantified in each run/replicate using intensities of detected signals. Runs were grouped for multivariate statistics on quantitative measurements to detect significant differential occupancy of p-sites. **Bioinformatic analysis:** Multivariate statistics on proteins and phosphopeptides were performed using Qlucore Omics Explorer 3.8 (Qlucore AB, Lund, *SWEDEN*) after a log2 transformation of abundance data for normalization. The transformed data were finally used for statistical analysis *i.e.,* evaluation of differentially present proteins/peptides between two groups using a Student’s bilateral *t* test and assuming equal variance between groups. Only proteins and peptides with a *p*-value < 0.05 were considered significant. Both volcano plot and heatmap views were generated with Qlucore Omics Explorer 3.8. Gene ontology analysis for cellular compartment using ShinyGo 0.77 (http://bioinformatics.sdstate.edu/go/) using a cutoff for fold change of ± 1.5 and *p*-value < 0.05. Metascape software (4) was used for the creation of protein association networks, with a cutoff of p<0.05. For Sequence Logo generation, all unique peptides with a fold change cutoff of ± 1.5 and *p* < 0.05 were completed through NCBI database searches to generate peptides of 15 amino acids lengths, with the phosphate in the middle. For Logo generation, the online platform for web logo was used [https://weblogo.berkeley.edu/logo.cgi](about:blank) (5).

**RNA extraction and real time qPCR**

Brain tissue from the hippocampus was homogenized and total RNA was extracted with TRIzol reagent (Invitrogen). Then, 1 µg of total RNA was reverse transcribed into cDNA following the High Capacity cDNA Reverse Transcriptase kit with random primers (Applied Biosystems). The quantification of relative mRNA levels was performed on a Realplex PCR machine (Eppendorf) using Taqman gene expression assays for CK2α (FAM Mm00786779_s1), and GAPDH (FAM Mm99999915_g1) as a housekeeping gene. Finally, relative mRNA expression was calculated with the ΔΔCt method (6).

**Hippocampal cell culture**

Hippocampal primary neurons from mice were cultured from P0 (post-natal day 0) mice. Brains were dissected in a dissection medium composed of HBSS (Gibco) and 10 mM Hepes. Hippocampi were collected, chemically dissociated with DNase (Roche) and Trypsin (Sigma) in a water-bath (20 min, 37 °C) and mechanically disaggregated. Following centrifugation (400 x g, 8 min) cells were counted and plated on 12-well plates coated with Poly-D-Lysine (Corning): 2.5 × 10^6^ cells/ml for Sholl analysis and 10 × 10^6^ cells/30 μl for immunocytochemistry. NeuroBasal medium supplemented (NB+) with 5% FBS (Biosera), B-27 (Gibco), GlutaMax (Gibco) and penicillin/streptomycin (Gibco) was used to grow the cells. To keep in the culture the glial cells at low levels at DIV1 half of the medium was replaced by NB+ with 2 μM AraC, and at DIV2 the medium replenished by a third of the final volume with NB+ with 4 μM AraC. Cell cultures were maintained at 37 °C with 5% CO2 until DIV14.

**Immunocytochemistry**

At DIV14, hippocampal primary neurons were fixed in 4% PFA/PBS for 10 minutes for Sholl analysis and in 4% PFA/4% sucrose/PBS for 20 min for immunocytochemistry. Following fixation, a permeabilization with PBS/0.5% Triton for 5 minutes was necessary. After three PBS rinses, coverslips were incubated with 3% BSA/PBS for 1 hr, RT. Cells were incubated with primary antibodies overnight at 4 °C. The next day, after three washes with PBS, they were incubated in the dark with secondary antibody. Coverslips were rinsed three times with PBS before mounting onto slides using Prolong^TM^ gold antifade reagent with DAPI (Invitrogen). Antibodies: chicken anti-MAP2 (ab5392, Abcam, 1:10000) and donkey anti-chicken Cy3 (703-165-155, Jackson ImmunoResearch, 1:400) for Sholl and mouse anti-PSD-95 (MABN68, Millipore, 1:500), guinea pig anti-VGLUT1 (135 304, SySy, 1:1000), rabbit anti-SynGAP1 (19739, Proteintech, 1:300), donkey A647 anti-mouse (A31571, Millipore, 1:200), goat A488 anti-guinea pig (GtxGp-003-F488NSX, ImmunoReagents, 1:400) and goat A488 anti-rabbit (A11008, Invitrogen, 1:400) for synapses. For Sholl analysis images were acquired with epifluorescence ZEISS Axioplan 2 microscope equipped with CoolSnap camera (Roper Scientific) using 20x/0.5 Dry objective and, 16-bit resolution of 1392 × 1040 pixels without digital zoom. For synapses, imaging was done with an inverted confocal microscope ZEISS LSM880 Elyra with Plan-Apochromat 63x/1.4 Oil DIC M27 objective and using confocal Airyscan mode of detection with digital zoom 1.8x. Images were acquired using bidirectional laser line average of 2 and with an 8-bit resolution of 1764 × 1764 pixels.

**Morphological analysis**

For analysis of dendritic arborization in hippocampal neuron cultures, a 2D Sholl analysis was performed with ImageJ (Sholl plugin). Images of isolated cells were thresholded in order to obtain binary images. Neuronal soma was selected, and the software counted the number of intersections by creating successive concentric circles with 5 μm step size from that area until 250 μm. The number of intersections was plotted as a function of the distance from the soma.

**Acute slice electrophysiology**

Transverse hippocampal slices were prepared from 4-5 months old Wt and heterozygous KI mice. Animals were anesthetized with ketamine (20 mg/kg), xylazine (1.4 mg/kg) and isoflurane, and perfused transcardially with a modified sucrose cutting solution containing the following (in mM): Sucrose 110, NaCl 10, KCl 2.5, NaH2PO4 1.25, NaHCO3 30, HEPES 20, glucose 25, thiourea 2, Na-ascorbate 5, Na-Pyruvate 3, CaCl2 0.5, MgCl2 10. Brains were rapidly removed, hippocampi dissected and placed upright into an agar mold and cut into 400 μm thick transverse slices (Leica VT1200S) in the same oxygenated cutting solution at 4 °C. Slices were transferred to an immersed-type chamber and maintained in carbogen (95% O_2_/5% CO_2_)-bubbled artificial cerebrospinal fluid (ACSF) containing the following (in mM): NaCl 125, KCl 2.5, NaH2PO4 1.25, NaHCO3 26, glucose 10, Na-pyruvate 2, CaCl2 2, MgCl2 1. Slices were incubated at 32 °C for approximately 20 min then maintained at room temperature for at least 1 h.

Prior to recording, slices were transferred to a recording chamber continuously perfused with warm (32.5 °C), oxygenated ACSF using a peristaltic pump, allowing perfusion with ACSF at 3 ml/min. The field recordings EPSPs (fEPSPs) were performed in current-clamp mode with a recording patch pipette (3–5 MΩ) containing 1 M of NaCl and positioned in the middle of stratum radiatum (SR) of hippocampal CA1 region. fEPSPs were evoked by mono-polar constant voltage stimulation (model DS2A, Digitimer) with a patch pipette filled with ACSF and positioned in CA1 SR. Pulses were delivered at 20-s intervals and stimulation voltage was set to obtain 30-40% of the maximum fEPSP. To induce LTP, we used a Theta-Burst Stimulation (TBS) protocol consisting of 5 bursts (5 pulses at 100 Hz per burst) interspaced by 200 ms repeated 4 times with a 2 second interval. Data were recorded with a Multiclamp 700B (Axon Instruments) and acquired with Clampex 10.7. The slope of the field EPSPs was measured using clampfit with all values normalized to a 20-min baseline period. All field recording experiments were done in the presence of gabazine (1 μM, SR-95531) and CGP56999A (2 μM), to block all GABA_A_ and GABA_B_ receptor mediated inhibitory transmission.

**Behavioral tests**

All behavioral testing was carried out in the light phase (light intensity: 45–50 lux). When repeatedly performed, such as fear conditioning, Barnes or Rotarod, experiments were performed at the same time on subsequent days.

**Y-maze: Spontaneous alternation test.** A Y-maze made of three black and opaque arms set at 120° angles was used. Visual cues were placed around the maze. The animal was placed at the center of the maze and was allowed to freely explore the three arms for 4 min. The number of arm entries and the order were annotated to calculate the percentage of alternation between the arms. An entry was considered when all four limbs of the animal were within the arm. An alternation is defined as successive entries in the different three arms on triple sets.

**Barnes Maze test**. Spatial learning was examined with the Barnes maze task that was performed as described in (7) with the following changes. The learning phase consisted of a training of 5 days (Days 1-5) with 3 sessions per day and inter-session time of 1 hour. After 2 days of rest, the reversal learning phase was performed for 4 days (Days 6-9) with 3 sessions per day. Test sessions were performed on sessions 3 of days 5 and 9, mice were allowed to explore for 1 minute without the escape box. Mice that did not explore were excluded of the analysis. For both learning phases the number of nose poke errors, the escape latency, the distance travelled and the velocity until finding the escape box were measured.

**Fear Conditioning.** Fear conditioning assesses the memory for the association between mild foot shock and a salient environmental cue. Freezing behavior, as defined as the complete lack of movement, is recorded since it is a characteristic fear response in rodents, in a hippocampal-dependent manner. Fear conditioning was performed in a testing chamber with internal dimensions of 25 × 25 × 25 cm^3^, which has transparent plastic walls each side and steel bars on the floor. The chamber was located inside a larger, insulated cabinet (67 × 53 × 55 cm^3^) to protect the test animals from outside noise. Training chambers were cleaned with 100% ethanol solution before and after each trial to avoid any olfactory cues. The experiments ran over 2 consecutive days. On Day 1, mice were placed in the conditioning chamber and 2 min 28 s later received one footstock (2 s, 0.3 mA). Mice were removed from the chamber 30 s after the shock. On Day 2, they returned to the same conditioning chamber for a 3-min period in the exact same conditions as Day 1 but without electrical shock, to evaluate context-induced freezing. Contextual fear conditioning experiment was carried out blindly with respect to genotype or treatment.

**Prepulse Inhibition (PPI)**. Mice were placed into acoustic startle chambers for a 40 min testing period. After the 5 min of acclimatization with permanent background noise (64 dB), four intensities of the prepulse were used: 72, 76, 80 and 84 dB. No-stimulus, prepulse or startle (120 dB) alone and prepulse-plus-pulse trials were pseudorandomly presented 10 times, with random intertrial interval times (10-20 s) separating the 20 ms long prepulse from the 40 ms long startle pulse during a total session time of 40 minutes. The intensity of the startle response was recorded in mV. PPI was calculated as a percentage of the pulse-alone startle amplitude. Startle amplitude was measured as the average voltage over the entire response window (mV-Avg).

**Overnight Recording.** Mice were singly housed in TSE systems metabolic cages ([PhenoMaster (tse-systems.com)](about:blank)) in the normal standard light-dark cycle, with free access to food and water for 48 hours. Locomotion, food and drink consumption were measured and analyzed with PhenoMaster software. **Rearing** during one hour (3-4 am the second night) was quantified while mice were in the metabolic cages.

**Nesting**. Animals were single-housed overnight, and three cotton rolls were added to each cage. After 16 hours nests were photographed, and scores were given following different criteria: 0 = Nestlet (cotton roll) remains nearly untouched (> 90% intact). 1 = Nestlet partly torn (50-90% intact). 2 = Nestlet mainly shredded but still 30-50% is not torn and the shredded ones are not gathered completely. 3 = Nestlet mostly shredded and gathered in one pad. 4 = Nestlet largely torn (> 90%), nest has a crater shape and at least 50% of its wall is higher than the mouse body.

**Marble burying test.** Anxiety and repetitive behavior were assessed twice, spaced by 7 days, by placing the mice in rat cages with 20 marbles arrayed on the surface of clean bedding. After 30 min the number of buried and not buried marbles was counted (they were considered buried when at least two-thirds of the surface was covered by the bedding).

**Open-Field (OF)**. Exploratory behavior and locomotor activity were measured in open-fields new environments (clear Plexiglas 40 x 40 x 40 cm) for 1 hour and data were collected with 10 min bins. The distance travelled was quantified by a computer-operated infrared photobeam activity system (Viewpoint version 5.31.0.120).

**Chemically-induced seizures**. Mice were injected with a single i.p. dose of picrotoxin (3 mg/kg in saline) and placed in cages in isolation for 60 min of observation. The time of onset of immobilization and first seizure, the number of seizures, and the total duration were annotated.

**Methods from supplemental text file:**

**Immunohistochemistry**

Perfused slices were washed in PBS and blocked in 3% goat serum (Jackson) in PBS/0.5% of Tween-20 (PBST) (1 hr, RT) followed by an overnight incubation with anti-CK2α (A300-198A, Bethyl, 1:1000). After three washes with PBST, slices were incubated with goat A594 anti-rabbit IgG antibody (A11037, Life Technologies, 1:400) for 1 hour at RT. After three rinses, slices were mounted with Prolong^TM^ Gold DAPI reagent (Invitrogen). Images were acquired with a ZEISS LSM880 Elyra with Plan-Apochromat 63x/1.4 Oil DIC M27 objective without digital zoom and using bidirectional laser line average of 2 and with an 8-bit resolution of 1912 × 1912 pixels.

**CK2 activity of extracts from mice brain tissue.** See methods for main figures.

**BRET**

HEK-293 cells were grown in DMEM supplemented with 10% (vol/vol) FBS, 1 g/L glucose, 1 mM glutamine. For BRET experiments, cells were transfected with the calcium phosphate precipitation method. Forty-eight hours after transfection, HEK-293 cells were washed twice with ice-cold PBS (Euromedex) and lysed in buffer containing 50 mM Tris pH 7.5, 150 mM NaCl, 10 mM EDTA, 1% Triton X100 and cocktail protease inhibitor (Roche) on ice for 10 min. Then, the lysates were centrifuged at 10 000 x g for 10 min. Laemmli buffer 4X (200 mM Tris-HCl pH 6.8, 4% SDS, 40% glycerol, 0.02% bromophenol and 0.5 M β-mercaptoethanol) was added to the supernatant. The cell lysates from HEK-293 transfected cells were separated by electrophoresis on SDS/PAGE (12% gels) and transferred to PVDF membranes (GE Healthcare Life Sciences). Blots containing YFP-tagged proteins were probed with a rabbit anti-GFP antibody (Cell Signaling, #2956, 1:1000). HRP-conjugated goat anti-rabbit IgG (H+L) antibodies (Invitrogen, #65-6120, 1:30000) were used as secondary antibodies. Immunoreactive bands were detected using the ECL detection kit (Biorad, 170-5060). with an Imaging System (Biorad, Chemidoc).

Forty-eight hours after transfection, HEK-293 cells were washed twice with ice-cold PBS (Euromedex) and resuspended in HBSS saline buffer (Invitrogen). Intact cells were distributed in 96-well microplates (Optiplate, Perkin Elmer). Furimazine substrate (Interchim B2W800) was added at a final concentration of 5 µM, and reading were performed with a Mithras LB 943 Multireader (Berthold), which allows the sequential integration of luminescence signals detected with two filter settings (NLuc filter, 480 ± 10 nm; YFP filter, 540 ± 20 nm). Emission signals at 540 nm were divided by emission signals at 480 nm. The BRET ratio was defined as the difference between the emission ratio obtained with co-transfected NLuc and YFP fusion proteins and that obtained with the NLuc fusion protein alone.

The results were expressed in milliBRET units (mBU, with 1 mBU corresponding to the BRET ratio values multiplied by 1000). BRET_max_ is the maximal BRET signal obtained in milliBRET units and BRET_50_ represents the ratio of acceptor and donor receptors (acceptor/donor) yielding 50% of maximum BRET signal. All BRET, luminescence, fluorescence measurements were performed at 21 °C using a Mithras LB 943 microplate analyser (Berthold). All data analyses were carried out with the GraphPad Prism 9.5.1 software for Windows (GraphPad Software Inc, San Diego, CA, USA). Concentration-response curves were fitted by nonlinear regression and saturation curves by a hyperbolic one-binding site equation. The method provided estimates for BRETmax and BRET50 values and corresponding SEM. Each experiment was repeated at least three times, and data were analyzed using one-way analysis of variance (ANOVA) followed by the Newman Keuls multiple comparisons test.

**Behavioral tests**

**Y-maze: Spatial recognition test.** During a 5 min habituation phase one random arm was closed by a door and the animal allowed to freely explore the Y maze, after which it was returned to its home cage. After one hour, the animal was tested during 4 min and the percentage of time the animal spent in the previously closed “novel” arm was measured.

**Barnes Maze test.** See methods for main figures.

**Overnight Recording.** See methods for main figures. **Rearing** for one hour (3-4 pm the second day) was quantified while mice were in the metabolic cages.

**Social interaction**. The **three-chamber test** assesses sociability in mice as they normally prefer to spend more time with another mouse then in an empty chamber. The test was carried out in 3-chamber boxes in Plexiglas (60 cm long, 40 cm large, 35 cm high). Boxes were divided into three compartments (chambers) of the same dimension by walls with small square openings (5 × 3 cm) allowing access into each chamber. Swiss mice of the same gender and age were used as “stimulus mice”. A video camera located at 1.5 m above the 3-chamber boxes was connected to an image analysis system (ViewPoint, France), allowing to record the distance travelled and the time spent by animals in the apparatus. One day before the experiment, mice were habituated to all chambers for 10 min. On the test day, the subject was confined in the center. Then, one empty cylinder and one cylinder with stranger 1 were placed in a corner of each side chamber. Doors were removed and the subject was allowed to explore the three chambers for 10 min after which it was re-confined to the center. Stimulus mouse 2 was placed in the previously empty cage, while stimulus mouse 1 was retained in its position. The subject was allowed to explore all arenas for 10 mins. Time spent exploring the boxes with or without the stimulus mice was recorded and the first 5 minutes plotted. The position of stimulus mice was randomized for all groups. **Quantification of nose pokes:** 2 mice of the same age, gender and genotype were introduced into an open field box (40 x 40 x 40 cm) and allowed to freely explore for 10 min. An observer blind to the genotype manually quantified the number or nose to nose and nose to body contacts between the pairs.

**Rotarod**. In order to examine motor function and balance of mice we used the accelerated rotarod test with settings: 4-40 rpm/5 min (Ugo Basile). The time that the mice were able to stay in the rod until fall was measured in seconds. After one training session, mice performed three consecutive trials per day with a 30 min inter-trial interval, for three consecutive days. If a mouse clinged on the rod for more than two full passive rotations, the mouse was removed from the trial and recorded as a fall.

**Grip Strength test.** To quantify muscular strength over time animals were tested from 5 weeks old to 15 weeks (3 trials, one day per week). Forelimb strength was measured with a digital grip strength meter (Bioseb). Mice were gently held by the base of the tail and allowed to grab the T-bar with the 2 frontal paws. With the torso in horizontal position mice were pulled back steadily until they released the bar.

**Inverted Screen Hanging test.** Mice were placed in the center of a wire grid which was then slowly inverted 180° and placed ~50 cm above a padded surface. The time until fall was measured, being 60 s the maximum hanging time.

**Light/Dark, novelty-suppressed feeding, tail suspension, forced swim and elevated plus maze tests** were performed as described (8).

**Fig. Legends Suppl figures**

**S1. Gross phenotyping of K198R(+/wt) mice.**

Lifespan of mice that had premature deaths according to sex and genotype. Percentages are calculated as the number of mice that died by the total number of born mice of that sex (A). Representative photograph of Wt and K198R(+/wt) male mice aged of 4 months (B). Food intake over 48 hours measured in metabolic monitoring cages showing no differences between genotypes (p = 0.5126, unpaired t test, n = 10 Wt and 13 Het) (C). All data are presented as mean ± SEM.

**Figure S2. No differences in subcellular CK2α localization in neurons.**

Representative images of coronal slices from Wt and K198R(+/wt) mice, probed with anti-CK2α antibody and imaged on a Zeiss LSM880.

**Figure S3. Optimization of parameters for intracellular CK2 activity assay**

Impact of total protein amount (A) and reaction time on CK2 activity (B). A reaction time of 15 min was used and was conducted with 90 mg of total protein per reaction. The final conditions of 30 min reaction time and 90 mg protein per reaction were selected for further experiments.

**Figure S4. Interaction of CK2α and CK2α using BRET.**

HEK-293 cells were transfected with CK2b-NanoLuc and either wt or of CK2α-YFP (R47G, K198R and R312W). Only upon interaction CK2α-β, a BRET signal at 530 nm is generated, as depicted in the scheme (A). BRET saturation curves were performed on HEK-293 cells transiently transfected with a constant DNA amount of CK2b-NanoLuc and increasing quantities of respectively either wt or of CK2α-YFP (R47G, K198R and R312W) (B). BRETmax quantification showed a reduction in R312W mutant (F(3,11) = 81.72, p <0.0001; 1-way ANOVA), but no difference for the other CK2α mutants (p = 0.9956 for R47G and p = 0.1632 for K198R; 1-way ANOVA, n = 3 independent experiments) (C). BRET50 which represents the ratio of acceptor and donor receptors (acceptor/donor) yielding 50% of maximum BRET signal showed no difference in mutants compared to wt (1-way ANOVA, n = 3 independent experiments) (D). All data are presented as mean ± SEM (*p < 0.05; **p < 0.01; ***p < 0.001; ****p < 0.0001). Measurements of CK2b-NLuc expression (luminescence) for BRET experiments. (A-D) Forty-eight hours after transfection, HEK-293 cells were detached with versene (Invitrogen) and resuspended in HBSS saline buffer (Invitrogen). Intact cells were distributed in 96-well microplates (Optiplate, Perkin Elmer). Furimazine substrate was added at a final concentration of 5 μM and total luminescence of cells was determined with a Mithras LB 943 Multireader (Berthold). Data are presented as mean ± SEM.

**Figure S5. Learning and memory paradigms.**

Y-maze recognition test shows no differences in the time that mice spent in the novel arm (p = 0.7474; Mann-Whitney test, n = 10 Wt and n = 14 Het) (A). Barnes maze test learning curves shows no differences between genotypes (n = 18 Wt and n = 14 K198R(+/wt)) for the number of error nose pokes (F(1,94) = 1.555, p = 0.2154 for Genotype) (B), nor the distance travelled to reach the target (F(1,94) = 0.2745, p = 0.6016 for Genotype) (C), but the escape latency was reduced in K198R(+/wt) (F(1,94) = 17.86, p < 0.0001 for Genotype) (D), and the velocity increased for Het compared to Wt mice (F(1,94) = 45.68, p < 0.0001 for Genotype) (E). Barnes maze test reversal learning curves shows no differences between genotypes (n = 18 Wt and 14 K198R(+/wt)) for the number of error nose pokes (F(1,93) = 0.5002, p = 0.4812 for Genotype) (F), nor the distance travelled to reach the target (F(1,93) = 0.2113, p = 0.6468 for Genotype) (G), but the escape latency is reduced in K198R/wt mice (F(1,93) = 20.47, p < 0.0001 for Genotype) (H). The velocity is increased in Het compared to Wt mice (F(1,93) = 14.19, p = 0.0003 for Genotype (I). For Barnes test, statistical analysis was performed using mixed ANOVA (Bonferroni multiple comparison tests) and data are presented as mean ± SEM (*p < 0.05; **p < 0.01;***p < 0.001; ****p < 0.0001).

**Figure S6. Various behavioral paradigms**.

Rearings recorded for one hour, at 3-4 pm on the second day of homecage activity recordings shows a reduction in K198R(+/wt) mice (p = 0.0411; unpaired t test, n = 20 mice per genotype) (A). Sociability was assessed in the 3-chamber test (F(1,112) = 1.211, p = 0.2735 for genotype; 2-way ANOVA; n = 16 Wt and n = 14 K198R(+/wt) (B), and by manual scoring of the number of sniffing/nose-to-nose pokes (p =0.6852, Mann-Whitney test, n = 10 Wt and n = 7 K198R(+/wt)) (C) or the duration of nose-to-nose pokes during a period of 10 min between two males of the same genotype during 10 min (p = 0.8868; Mann-Whitney test) (D). In the rotarod test, the latency of K198R/wt and control mice to fall off the rod during accelerated shows no difference between genotypes (F(1,33) = 2.789, p = 0.1044 for Genotype, 2-way ANOVA, Bonferroni multiple comparison tests, n = 18 Wt and n = 17 K198R(+/wt)) (E). Mice strength was assessed in the grip test and shows no differences between genotypes (F(1,20) =0.3351, p = 0.5691 for Genotype, mixed ANOVA, n = 14 Wt and n = 8 Het) (F). Inverted screen test shows no differences in the time hanging between genotypes (p = n 0.2107; unpaired t test, n = 13 Wt and n = 12 Het) (G). All data are presented as mean ± SEM (*p < 0.05; **p < 0.01; ***p < 0.001; ****p < 0.0001).

**Figure S7. Mood assessing behavioral paradigms.**

Light/Dark transition test shows that compared to Wt, K198R(+/wt) mice show no preference for the dark compartment (p < 0.0001 and 0.8470, respectively; Mann-Whitney test, n = 12 Wt and n = 11 Het) (A). Novelty suppressed feeding shows reduced latency to feed in Het (p = 0.0369, unpaired t test, n = 8 Wt and n = 6 Het) (B).

Elevated plus maze shows no differences for the different arms between genotypes (F(1,69) = 0.001125, p = 0.9733 for Genotype, 2-way ANOVA, Bonferroni multiple comparison tests, n = 12 Wt and n = 13 Het) (C). Tail suspension (D) and forced swim test (E) show no difference between genotypes (p = 0.1932 and p = 0.0766, respectively, Mann-Whitney, n = 12 Wt and n = 14 Het). All data are presented as mean ± SEM (*p < 0.05; **p < 0.01; ***p < 0.001; ****p < 0.0001).
